# PATHOGEN GROWTH AND VIRULENCE DYNAMICS DRIVE THE HOST EVOLUTION AGAINST COINFECTIONS

**DOI:** 10.1101/2024.05.01.592035

**Authors:** Srijan Seal, Dipendra Nath Basu, Kripanjali Ghosh, Aryan Ramachandran, Rintu Kutum, Triveni Shelke, Ishaan Gupta, Imroze Khan

**Affiliations:** Trivedi School of Biosciences, Ashoka University, Sonepat, Haryana 131029, India; Department of Biochemical Engineering and Biotechnology, Indian Institute of Technology, Hauz Khas, New Delhi-110016, India

**Keywords:** Adaptive dynamics, Coinfection, Experimental evolution, Innate immunity, Transcriptomics

## Abstract

Coinfections, or the simultaneous infection of hosts by multiple pathogens, are widespread in nature with significant negative impacts on global health. Can hosts evolve against such coinfections as effectively as they would against individual pathogens? Also, what roles do individual pathogens play during such evolution? Here, we combined theoretical models and experiments with *Tribolium castaneum* populations evolving against two coinfecting bacterial pathogens, with contrasting growth and virulence dynamics, to reveal that fast-growing pathogens inflicting rapid mortality surges (i.e., fast-acting) restrict adaptive success against coinfections. While hosts rapidly evolved better survival against slow-growing bacteria causing long-lasting infections, evolution against coinfection was significantly delayed and resembled slow adaptation against fast-acting pathogens. Moreover, limited scopes of immunomodulation against fast-acting pathogens during coinfections can drive the observed adaptive patterns. Overall, we provide new insights into how adaptive dynamics and mechanistic bases against coinfections are critically regulated by individual pathogens’ growth and virulence dynamics.

## INTRODUCTION

Coinfection of a host by multiple pathogen species is highly ubiquitous (*1–3*). Although biomedical research has primarily focused on isolated interactions of a single host vs. a single pathogen, growing evidence from natural systems and epidemiological studies indicates the greater ecological importance of co-infecting pathogens in influencing the global health and disease burden (*4*, *5*). For example, in humans, over one-sixth of the world’s population is affected by coinfections, comprised of diverse pathogens underlying many globally important diseases such as HIV, tuberculosis, malaria, hepatitis, and leishmaniasis (*6–8*). In addition to engaging in complex within-host interspecific interactions (*1*), the evolutionary history of frequent exposure to these coinfecting pathogens can profoundly influence the maintenance and deployment of the host immune system (*9*, *10*). However, despite such natural relevance, whether or to what extent coinfecting pathogens could influence the evolution of the host immune system differently from infections caused by single pathogens remains unexplored.

A generalized understanding of coinfection outcome is also challenging because of multiple confounding parameters such as the multiplication rate of individual pathogens during coinfection (*11*), temporal changes in their relative frequency and damage to the host (*12*) that influence the dynamics and efficacy of the immune activation. For instance, pathogens with divergent antigenic properties, which a single immune strategy cannot control, might lead the host to activate multiple immune components simultaneously during coinfection, increasing the energetic burden (*13*) and immunopathological risk (*14*, *15*). Moreover, many naturally occurring coinfection can also involve pathogens that vary widely in their growth and virulence dynamics (*16*). In such cases, the host might evolve temporally separated immune strategies depending on how and at what rate different pathogens multiply inside the host and manifest their virulence (*17*). For instance, the immune system might experience a strong selection to rapidly eliminate the fast-growing pathogens that induce high mortality rates early in infection (i.e., fast-acting) (*18*, *19*). Recent experiments and theoretical models can support this idea, where the ability to effectively clear pathogens by mounting appropriate immune responses early in the infection can serve as a critical determinant of post-infection survival success (*20*, *21*). However, the evolution of host responses facilitating such early-life fitness advantages against coinfections can be constrained if the response time to fast-acting infections is limited, thereby precluding the timely induction of appropriate immune components at adequate levels (*22*, *23*). This can be further complicated by the co-occurrence of other pathogens that grow relatively slowly, induce a slower mortality rate and persist longer (i.e., slow-acting), thereby warranting a sustained immune activation (*24*, *25*). Consequently, the efficacy of immune adaptation against coinfections might be critically contingent upon balancing the expression of such specific immunomodulation against individual pathogens (*25*, *26*). However, experiments accounting for the differences in growth and virulence dynamics between co-infecting pathogens while analysing their impacts on immune system evolution are missing.

To fill these gaps, we first built a theoretical model that describes diverse adaptive trajectories of host responses against coinfections based on their differences in within-host pathogen growth rate, rate of clearance, timing of immune activation, host mortality rate (i.e., virulence manifestation), and the level of interference due to competitive interactions and immune cross-reactivity between pathogens (*27*, *28*). Our model predicts that for pathogens that do not strongly interfere with each other’s growth or induce strong cross-reactive immunity, within-host growth dynamics of rapidly proliferating pathogens and their effects on host mortality rate at the early infection phase can drive the adaptive dynamics against coinfections. Subsequently, we tested this prediction using experimentally evolving *Tribolium castaneum* beetles adapting against two coinfecting bacterial pathogens with contrasting growth and virulence dynamics for 30 successive generations. At every generation, beetles were infected with either (A) fast-growing Gram-positive bacteria *Bacillus thuringiensis* (Bt), causing a rapid and sharp increase in host mortality followed by rapid clearance within a day (i.e., fast-acting) or (B) slow-growing Gram-negative bacteria *Pseudomonas entomophila* (Pe) that killed the beetles at a slower rate, while causing persistent infection for several weeks (i.e., slow-acting); or (C) a combination of both the pathogens (Mx) (see **Fig. 1** for study design). Finally, we used an RNA-sequencing approach to investigate the underlying changes in gene expression to gain comparative molecular insights into host adaptations against individual vs coinfecting pathogens.

**Figure 1:**
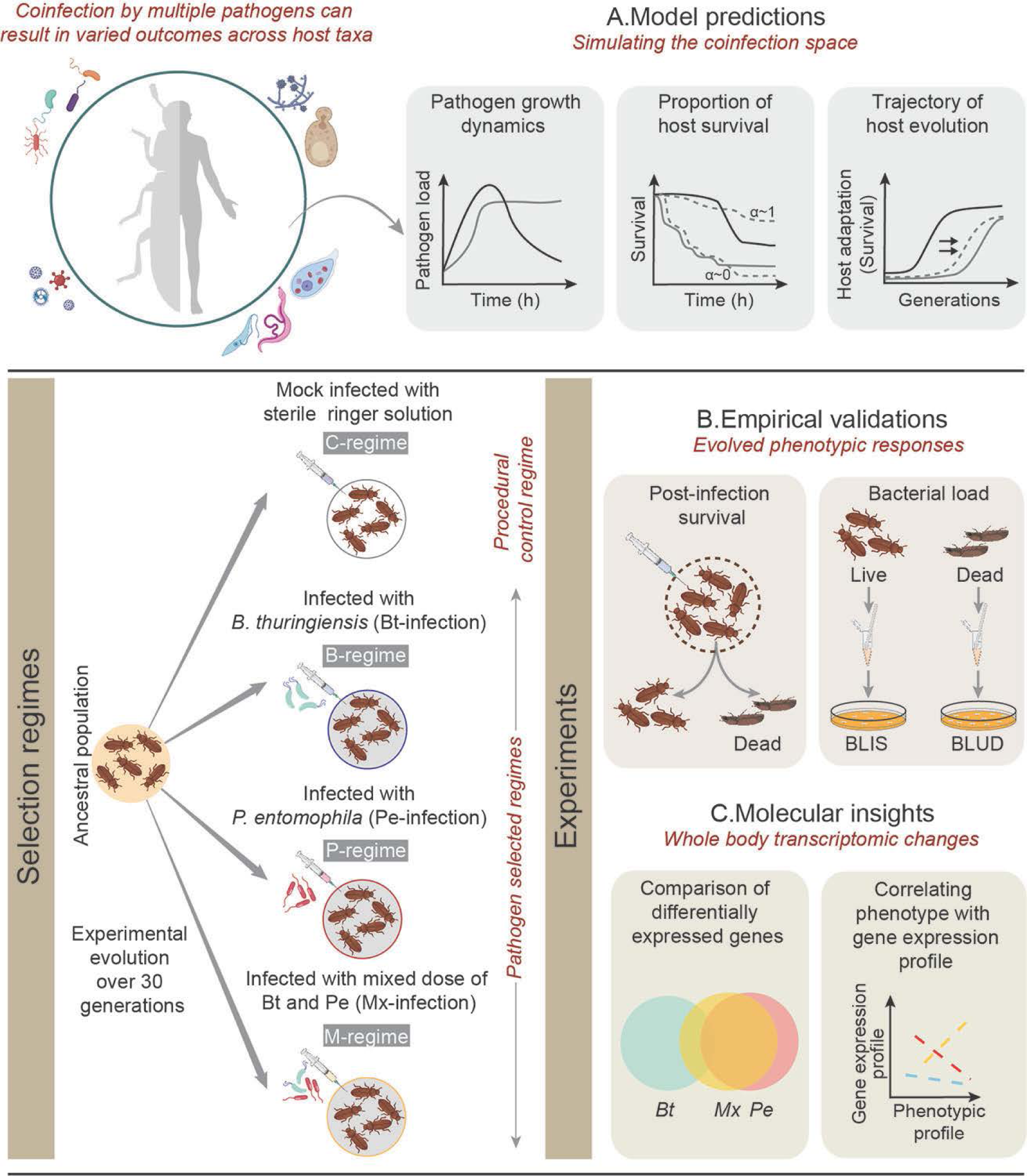
A brief description of study design. **(A)** First, theoretical modelling to predict the adaptive dynamics against coinfections (dotted line) based on the within-host growth vs clearance rate of individual pathogens (solid line), timing of immune activation, the host mortality rate caused by each pathogen and their level of interference (*α*) ; **(B)** Subsequently, empirical validation of the predicted patterns of adaptive dynamics against coinfection, using experimentally evolving model insect *Tribolium castaneum* beetles adapting against two coinfecting bacterial pathogens with contrasting growth and virulence dynamics. Replicated beetle populations were infected with either fast-growing *Bacillus thuringiensis* (Bt) or slow-growing Gram-negative bacteria *Pseudomonas entomophila* (Pe) or a combination of both the pathogens (Mx) to create three pathogen-selected regimes such as B-, P- and M-regimes, respectively (n= 4 independent replicate populations with 75 breeding pairs/ regime). We also had unselected control populations (C-regime) where beetles were sham-infected with sterile insect Ringer solution. We used post-infection survival and reduction in **<u>b</u>**acterial **l**oad **i**n **<u>s</u>**urviving (BLIS) beetles as proxies for the adaptive evolution of pathogen resistance, whereas **<u>b</u>**acterial **l**oad **<u>u</u>**pon **<u>d</u>**eath (BLUD) served as a measure of lethal pathogen burden across selection regimes; **(C)** Finally, we used an RNA-sequencing approach to analyse underlying changes in gene expression profiles to gain molecular insights and explain the observed phenotypic variations during the experimental evolution.

We speculated two alternative possibilities— (A) If variations in the beetle’s ability to clear fast-growing Bt primarily determine their survival probability early in the coinfection (*20*, *29*), selection pressure might act more strongly to resist the infection prevalence of Bt than Pe. Consequently, the rate of adaptation against Mx might closely resemble the responses against only Bt infection.

Moreover, there could also be a delay in their rate of adaptation if the scope of immune modulations is limited against the early mortality surges caused by rapid-acting Bt (*30*, *31*);(B) Alternatively, since Bt-induced early infection phase is closely followed by a persistent Pe infection phase that interferes with the beetle’s oviposition window, selection can instead be stronger against long-lasting Pe infections to ameliorate its fitness costs during reproduction (*32*). This could bias the overall adaptive dynamics more towards the responses against Pe. Our results supported the model outputs such that fast-acting pathogens such as Bt imposing early infection costs indeed constrained the adaptation against Mx. Mechanistically, the observed patterns of adaptation against Mx could result from fewer immunomodulatory mechanisms available against its fast-growing Bt counterparts. Overall, these are rare insights into how selection against individual pathogens can determine the trajectory of phenotypic variations vs mechanistic changes while evolving against coinfections.

## RESULTS

### The theoretical model predicts the importance of within-host pathogen growth and virulence dynamics in understanding the host adaptation against coinfections

To understand the host adaptation against coinfecting pathogens with contrasting growth dynamics, we began by first simulating the density-dependent growth rate of individual pathogens (e.g., rapid vs slow) leading to the acute infection phase using a Baranyi model, as described in Duneau et al. (*20*). Here, we also considered varying initial inoculation sizes, the lag phase of pathogen growth, and carrying capacity across pathogens and infection types. In this model, a subset of individuals succumbed to infection due to their inability to control the pathogen growth below a threshold density, causing terminal infection. Following this, we also simulated the divergent clearance patterns of the pathogens from the surviving individuals, leading to either rapid clearance or long-lasting persistent infection (i.e., incomplete clearance) using an exponential decline model (*20*) (See **Table S1** for parameters). Overall, this enabled us to typify within-host growth dynamics patterns of pathogens that could constitute diverse facets of a two-pathogen coinfection system: e.g., rapidly growing pathogens causing acute infections, followed by rapid clearance (Rc) or persistent infection (Rp); Slow-growing pathogens causing acute infection, followed by rapid clearance (Sc) or persistent infection (Sp) (**Fig. 2A**). Subsequently, we paired these pathogens based on their contrasting growth rates (i.e., rapid vs slow) to create the following coinfection combinations: e.g., Rc-Sc, Rc-Sp, Rp-Sc and Rp-Sp. Note that we also considered a coefficient of interference α, describing the effects of direct competitive interactions (*β*) between co-infecting pathogens (*27*) and cross-reactive immune modulations (*γ*) affecting their growth (*28*), which increases when either of the coexisting pathogens attains their peak growth (**Fig. 2B**). Moreover, in the case of pathogens that can be rapidly cleared by the host, the level of their interference can also decline rapidly, whereas pathogens causing persistent infections, can retain their interference longer as they enter chronic infection phase (**Fig. 2B**). Nevertheless, growth dynamics, as well as the nature of interference between pathogens, jointly influence the host survival. In the absence of strong interference (i.e., low *β*- and *γ*-values), the early host mortality pattern due to coinfection closely resembles the early host mortality trend against the rapid-acting pathogen that grows and imposes mortality rapidly (**Fig. 2C, Table S2**). By contrast, pathogens with very high mutual interference can lead to less severe effects of their coinfection relative to their single infection counterparts.

**Figure 2:**
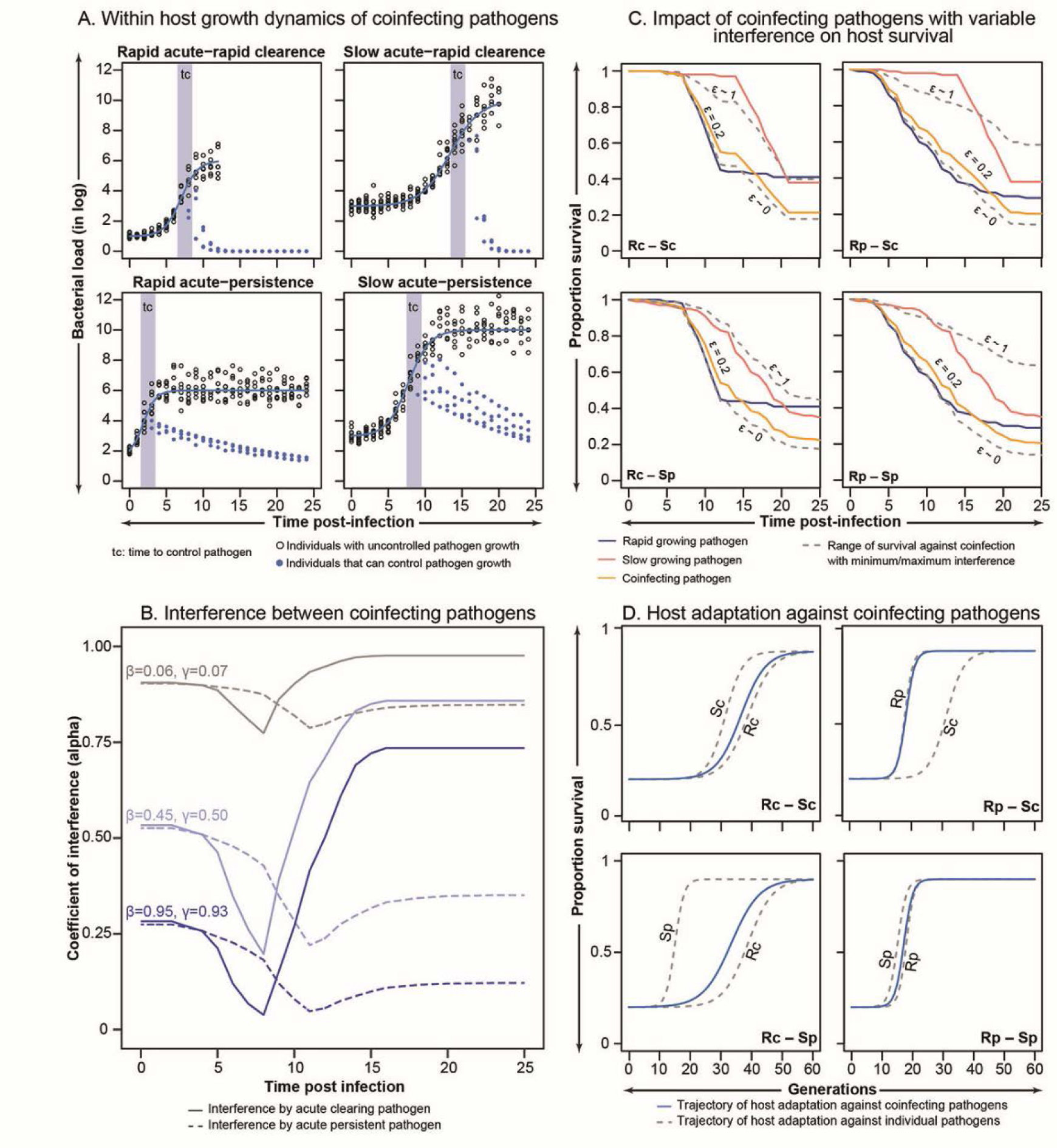
A mathematical model of coinfection landscape and host evolution. **(A)** Combination of pathogens with divergent growth dynamics during coinfection: fast-growing pathogens causing acute infections, followed by (i) rapid clearance (Rc) or (ii) persistent infection (Rp); Slow-growing pathogens causing acute infection, followed by (iii) rapid clearance (Sc) or (iv) persistent infection (Sp), using the combination of Baranyi model and an exponential decline model, developed by Duneau et al. 2017 (n=250; 10 data points for each of the 25 time points); **(B)** The shape of the coefficient of interference (*α*) was simulated for both rapidly cleared vs persistent pathogens, as well as for the various extent of *β* and *γ*; The coefficient of interference α between coinfecting pathogens is dependent on pathogen growth dynamics, and the parameter *ε* (combining the interference due to host immune modulations by the co-infecting counterpart (*β*) and direct resource-driven competition between the co-infecting pathogens (*γ*)); **(C)** Probable effects of coinfection by different combinations of pathogens (as described in **A**) on host survival, based on survival patterns against individual pathogens by incorporating conditional probabilities of survival and variable degrees of interference between pathogen types (as described in **B**); **(D)** The host adaptive trajectories across various combinations of rapid-vs slow-growing pathogens only at low *ε* values. The trajectories were determined by the growth dynamics of rapidly proliferating pathogens and their total infection window.

Since low interference between pathogens increases the severity of coinfections (estimated as post-infection mortality described above (**Fig. 2C**), we assumed this to be the most relevant condition that maximizes the selection on the host to reduce the infection costs. We thus only modelled the host adaptive trajectories across various combinations of rapid-vs slow-growing pathogens only at low *β*- and *γ*-values (**Table S3**). Overall, our model predicts that the rate of host adaptation against coinfection is determined by the growth dynamics of rapidly proliferating pathogens and their total infection window (i.e., from the first pathogen exposure to the end of mortality due to infection), where host mortality happens, and fitness costs can be paid under pathogenic infections (**Fig. 2D, Table S3**), overriding the effects of growth and virulence dynamics of slow-proliferating pathogens. For example, rapidly growing pathogens such as Rc, which can lead to early mortality surges within a short time, can significantly delay the host adaptation against Rc-Sp or Rc-Sc combinations. In contrast, host evolution (i.e., gain in the survival advantage against pathogens) was faster when the rapid growth phase was followed by persistent infection, causing mortality over a prolonged period (i.e., Rp-Sp and Rp-Sp).

### Experimental data confirms that rapidly growing pathogen determines the coinfection outcome during the early infection phase

We next performed a series of experiments to verify the above model predictions, underscoring the role of rapidly growing pathogens in driving the evolution against coinfections. We chose to verify the host adaptive trajectory against a pair of coinfecting pathogens, which are already predicted by the model to produce the most contrasting effects on host adaptation attributed to their divergent growth and virulence dynamics— i.e., Rc-like pathogen in combination with another slow-growing and slow-killing Sp-like pathogen (**Fig 2D**). To this end, we used the model insect *T. castaneum* infected with a mix of suitable bacterial pathogens, *B. thuringiensis* (Bt) and *Pseudomonas entomophila* (Pe), which we identified as possessing comparable growth and virulence dynamics as that of Rc- and Sp-like pathogens, respectively. Both pathogens killed ∼60–65% of the beetles within a week, although the mortality rates differed. While Bt-induced mortality showed a rapid surge within 8 hours post-infection (hpi), with most susceptible individuals dying within the first 12hpi (**Fig. 3A; Table S4**), Pe-induced mortality showed a relatively late onset of around 20–24hpi and continued for the next 7-days. However, beetles infected with a combination of Bt and Pe (Mx) showed a mixed mortality pattern reflecting the individual effects of both pathogens such that there was a sharp Bt-like decline in their survival within the first 16–18hpi, which is then followed by a gradual Pe-like decline for the next 7 days (**Fig. 3A**). During this, Bt cells showed rapid growth between the first 6–8hpi and then became undetectable by 20 hours, whereas the Pe cells reached peak growth around 24hpi and persisted in high numbers even after a week (**Fig. 3B & C; Table S5**) (Pe cells persisted even after 25 days post-infection in some beetles; **Fig. S1**). Moreover, in beetles infected with Mx, early mortalities (within 20hpi) were primarily driven by Bt-induced pathogenicity as dead beetles carried a large abundance of Bt cells (∼10^6^ cells/ beetle; estimated immediately after death), whereas later mortalities (>20hpi) were most possibly caused by an overgrowth of only Pe (∼10^7^ cells/ beetle) (**Fig. 3D**). These results thus corroborate the model predictions where the severity of coinfection and host mortality patterns during early infection phase was indeed correlated with the effects of rapid-growing pathogen counterparts (See **Fig 2C**).

**Figure 3:**
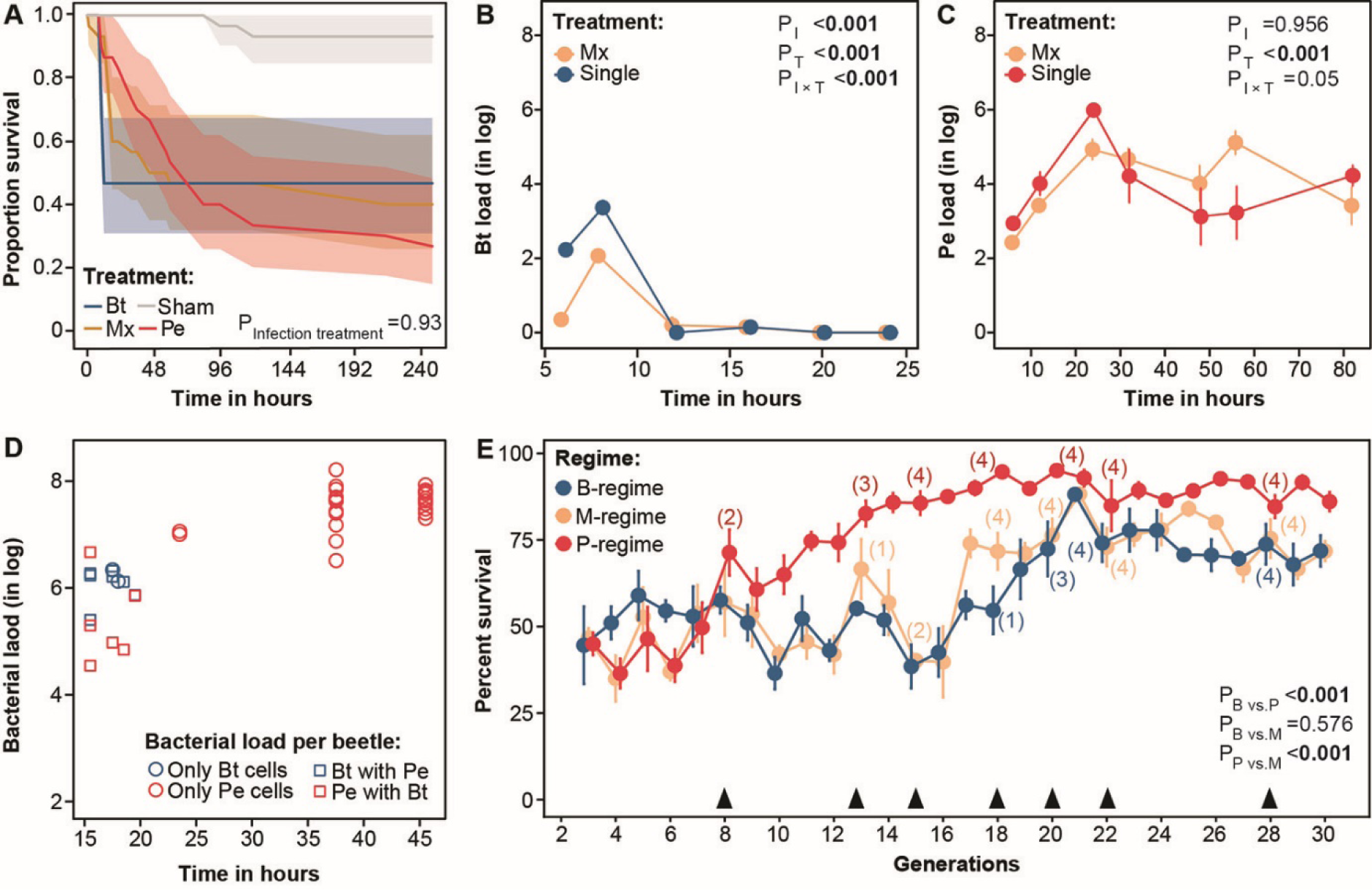
**(A)** Proportion of beetles from baseline population (henceforth, baseline beetles) surviving after infection with bacterial pathogens *B. thuringiensis* (Bt), *P. entomophila* (Pe) or a combination of both (Mx) (n= 30 females/treatment). The P-value represents differences across different infection treatments (i.e., Bt, Pe, and Mx); Temporal changes in within-host growth dynamics of **(B)** Bt and **(C)** Pe load in baseline beetles, both in the context of infections caused by single- (i.e., Bt or Pe) vs co-infecting pathogens (i.e., Mx) (n= 10 replicates with pooled homogenate of 3 females/ timepoints/infection treatment). P-values represent the effects of infection treatment (I) and the assay time (T); **(D)** The load of Bt vs Pe cells in baseline beetles that succumbed to infection, assayed till first 46h after Mx infection. Bt cells were detected only in beetles that died within the first 19h of infection (n=9). At later time points (>19–45h), beetles (n=25) only carried Pe cells; **(E)** Beetle survival across generations (Generation 3–30) at the end of the oviposition window (i.e., 8th-day post-infection) in each replicate population of different pathogen-selection regimes (n= 4 replicate populations/selection regime). P-values represent the pairwise differences between selection regimes. Solid black triangles denote the generations where selection response was assayed by comparing the post-infection survival of control (C) vs pathogen-selected regimes (B, P and M) against their respective pathogens (n=24–60 females/regime/generation). The numbers in the parentheses represent the number of replicate populations from each selection regime that showed significantly improved survival during experimental evolution (also see **Fig. S2**). All the assays involved 4 replicate populations, except generation 8, where only 3 replicate populations could be assayed.

### Experimental data validates that rapidly growing pathogen drives the host adaptive dynamics against coinfection

We next allowed beetle populations to evolve under strong pathogen selection imposed by either Bt (B-regime) or Pe (P-regime) or a mix of both (M-regime), each with 4 replicate populations (i.e., B1– 4; P1–4; M1–4) and tracked their post-infection survival for 30 generations. Beetle response against Pe was the fastest so that within only 8 generations, they could rapidly increase their post-infection survival from ∼40 to ∼75% and then to ∼90% by 18 generations (**Fig. 2E; Table S6;** also see **Fig. S2).** In contrast, B and M beetles required a substantially extended selection period to improve survival. They initially showed large fluctuations in survival (∼35–60%) for 16 generations and then could steadily increase only up to ∼75% by the 24^th^ generation. Control populations that were either pricked with sterile Ringer solution (C) (or maintained as unhandled populations) had a very high survival rate (>98%) throughout the experiment. Parallelly, we also directly estimated the relative improvement in post-infection survival of each replicate population, relative to C beetles, at regular intervals between generations 8–28 to disentangle the adaptive dynamics across pathogens and infection types (**Fig. 2E, S3)**. While at least half of the replicate P-populations showed significantly improved survival within 8 generations, followed by the other two populations by 15 generations, the first replicate population of M- and B-beetles could evolve the response only at generations 13 and 18, respectively. The remaining M- and B-populations evolved the response after the 18^th^ and 22^nd^ generations. Overall, while these results highlight the divergence in the rate of adaptation across pathogens (e.g., Bt vs Pe) and infection types (single vs multiple pathogens), they are also in conformity with the theoretical predictions, emphasizing the role of fast-growing Bt-like pathogens in restricting the adaptation against coinfections.

We also found a significant reduction in the bacterial load across pathogen-selected regimes relative to C-beetles, estimated around the onset of mortality after Pe and Bt infections (i.e., 24 and 8hpi, respectively) and at two-time points after Mx infection (8hpi and 20hpi) (**Fig. S4, S5; Table S7–9**). Increased post-infection survival of evolved beetles could thus be associated with their improved ability to prevent bacterial growth relative to the unselected control beetles.

### Host populations evolving against coinfection adopted distinct strategies to counter the severity of infections caused by individual pathogens

Next, we also compared the bacterial load of every M- and C-beetle that succumbed to Mx infection and sampled a subset of survivors every 5–8h for the next 50hpi to explain their divergent mortality patterns as a function of temporal changes in the pathogen growth dynamics. Contrary to our expectation, live M- beetles did not carry fewer Bt cells than C-beetles, except during the early phase of infection before mortality was initiated (i.e., 6–8 hours) (**Fig. 4A; Table S10, also see Fig. S4F**). However, they could significantly limit the number of beetle mortalities due to the growth of Bt cells beyond a threshold density at the early phase of infection (i.e., compare the number of dead beetles in M- and C-regime between 12–20hpi; **Fig. S6; Tables S11**). Interestingly, most of the live M- and C- beetles showed complete removal of Bt cells within ∼20 hours of infection (**Fig. 4A**), suggesting no differences in their rate of pathogen clearance (**Fig. S7; Tables S12**). Increased efficacy in arresting the Bt growth below the threshold density causing terminal infection, rather than its clearance, thus explained the improved survival of M-beetles relative to C-beetles.

**Figure 4:**
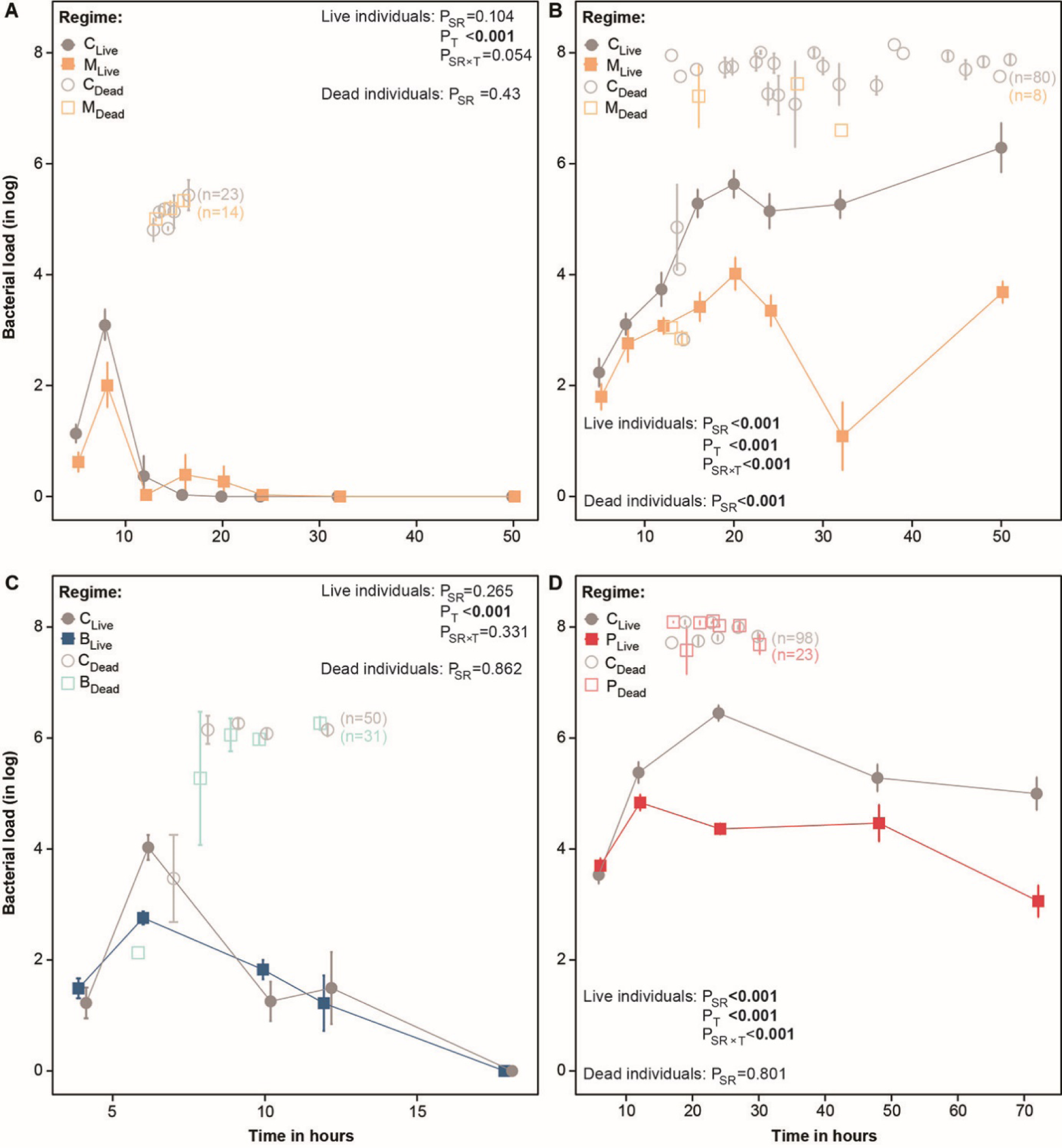
Within-host bacterial growth dynamics in control vs selected beetles. Temporal changes in **(A)** *B. thuringiensis* (Bt) **(B)** *P. entomophila* (Pe) load of live (n= 6–10 replicates with pooled homogenate of 3 females/selection regime/time point/ bacteria) and dead beetles sampled at various time points after coinfection in M regime relative to C regime until 50 hours post-infection (hpi); Temporal changes in **(C)** Bt and **(D)** Pe load sampled from live (n= 6–10 replicates with pooled homogenate of 3 females/selection regime/time point/bacteria) and dead beetles in B and P regime, relative to their C counterparts. Bt load from every dead B vs C beetle was recorded till 12hpi, whereas live individuals were monitored till 18hpi as Bt cells are usually cleared by beetles by this time. In contrast, Pe load from dead and live P vs C beetles was recorded only till 30hpi (beetle mortality beyond this point was not tracked for bacterial load assay) and 72hpi, respectively. In each case, bacterial load of dead beetles was extracted from individual beetles. The total number of beetles that died is indicated in parentheses. In each panel, P-values either represent the effects of the selection regime (SR) and assay time (T) on bacterial load derived from live individuals; or the main effect of the SR on bacterial load upon death.

We noted that the number of M-beetles that died due to Pe overgrowth after 16h was also drastically reduced (**Fig. 3B; Table S10**), but now, in contrast to the Bt-infection phase, surviving M beetles always carried a much lower density of Pe cells (**Fig. 3B; Table S10**), indicating that evolved beetles cleared Pe more efficiently. This, in turn, enabled them to prevent the Pe load of surviving beetles from exceeding the threshold density, leading to lethally acute infection (**Fig. 3B)**. Also, M- and C-beetles that succumbed to infection did not differ in their Bt or Pe burden, suggesting that the threshold pathogen density needed to cause mortality was comparable across regimes (**Fig. 3A, B; Table S10**). Overall, these results broadly corroborated the patterns of bacterial growth dynamics in B- and P-regimes as well (**Fig. 3C, D; Table S10**), suggesting that the outcome of Mx infection in the M-regime might be additively determined by both their initial success in controlling the Bt overgrowth as well as maintaining lower Pe burden in the later phase of infection.

### Immune gene expression profiles in host populations adapted against the coinfection resembled more with those evolving against the slow-growing pathogen

To gain mechanistic insights into divergent responses evolving across pathogens and infection types, we next conducted RNAseq using beetles across selection regimes, collected around the onset of their mortality after respective infection treatments (i.e., 8, 16 and 24h after infection with Bt, Mx, and Pe, respectively). This allowed us to compare the gene expression changes underlying nearly comparable fitness consequences across diverse beetle lines and infection types. Overall, the number of differentially expressed genes (DEGs) upon infection was considerably higher in M- beetles and P-beetles compared to B-beetles, both before (No. of genes: M= 427, P= 439, B= 165) and after (No. of genes: M= 374, P= 472, B= 171) the experimental evolution (**Fig. S8A, B**). Also, the evolved M-beetles showed a significantly higher number of overlapping DEGs with that of P-beetles (N= 119) than B-beetles (N=29), which might indicate similar mechanisms using a shared set of candidate genes between M- and P-beetles (**Fig. S8C**). We found 77 and 81 DEGs common across infection treatments in control and evolved populations, respectively. Those common set of genes possibly played pervasive roles across pathogens and infection types including immune-related molecules such as peptidoglycan recognition proteins (PGRP SC2), gram-negative bacteria binding proteins, antimicrobial peptides (AMPs; Attacin 2, Coleoptericin and Defensin 3) as well as key metabolic genes namely glucose dehydrogenase and fatty acyl CoA reductase. We also identified 65 DEGs upon infection with known immune functions across pathogens and selection regimes (**Table S13**). However, to disentangle their roles, we divided them into five broad categories based on their immune-related functions (i.e. immune categories) (*33–35*): (a) pathogen and immune receptors; (b) immune regulators; (c) inducible immune effectors, including AMPs and lysozymes; fast-acting constitutively expressed (d) melanisation response involving phenoloxidase pathway; and (e) production of reactive oxygen species (**Fig. 5A**), followed by a MANOVA to test effects of infection status, pathogen identity and selection regimes on each of these immune categories (**Tables S14– S18**). While the effects of selection regimes varied across immune categories, the infection status and pathogen identity produced the most consistent changes. They also showed a strong two-way interaction across immune categories, suggesting the deployment of pathogen-specific immune responses.

**Figure 5:**
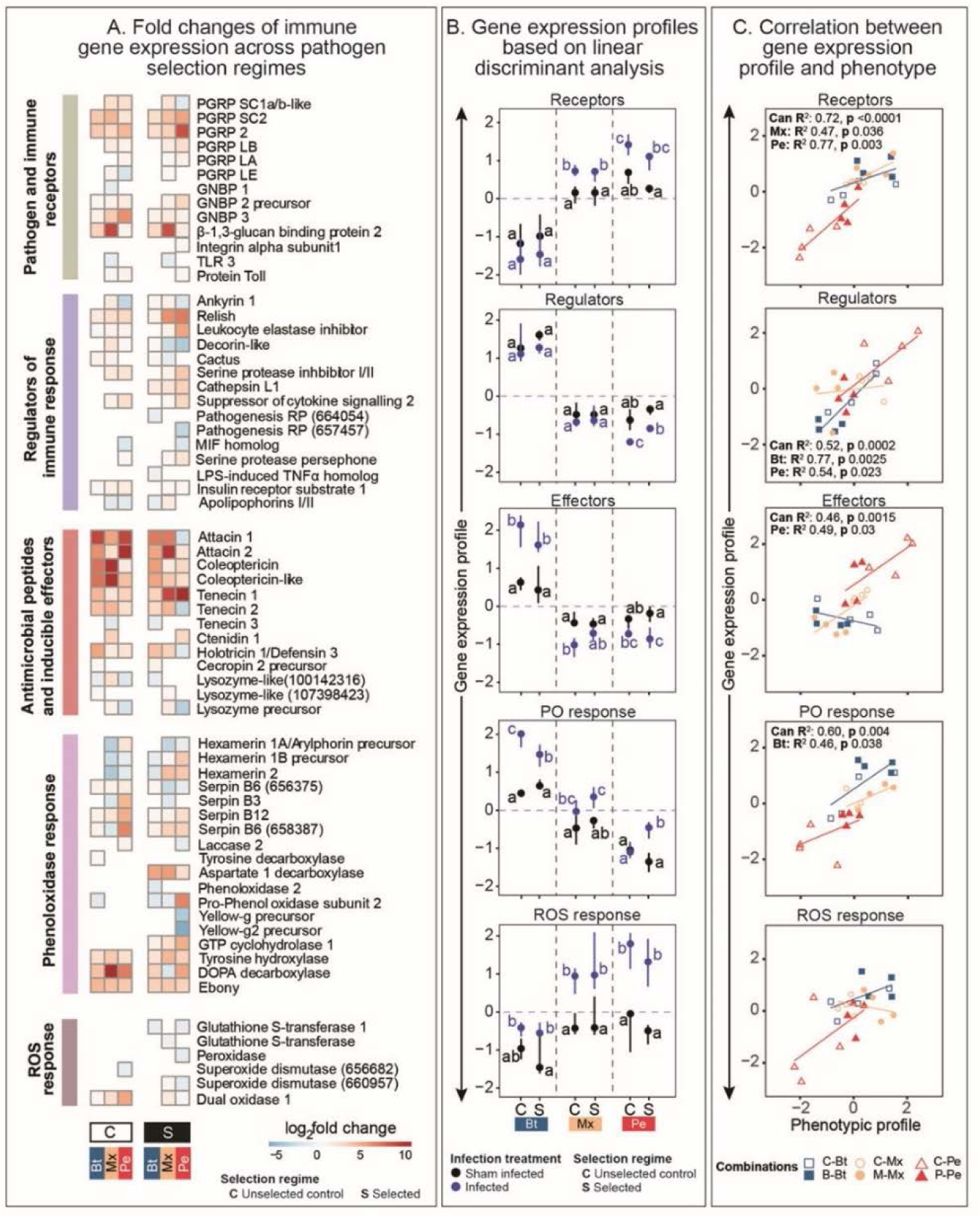
RNA sequencing and molecular insights into the evolved responses. **(A)** Heatmaps denoting the differentially regulated genes with known immunological function in insects (described in **Table S14**) after respective infection treatments (e.g., sham infection vs either Bt, Pe, or Mx) in control vs pathogen-selected regimes (B-, P- or M-regimes), divided into five broad functional categories: (a) pathogen and immune receptors; (b) immune regulators; (c) inducible immune effectors, including antimicrobial peptides (AMPs) and lysozymes; (d) melanisation response involving phenoloxidase pathway; and (e) production of reactive oxygen species; **(B)** Cumulative gene expression profile of differentially expressed immune genes based on linear discriminant analysis (LDA). The first axis of LDA is considered as gene expression profile for various categories of differentially expressed immune-related genes. Here, we compared expression profile changes between beetles with sham infection and infection with respective pathogens (Bt, Pe, and Mx) in pathogen-selected (S) beetles (B-, P-, M-beetles) vs their respective unselected (C) beetle populations. In each panel, significantly different groups are connected with different alphabets. Alphabet assignments are not comparable across pathogens (separated by dotted lines); **(C)** Correlation of phenotypic profile (based on combined estimates of post-infection survival and bacterial load) with cumulative expression levels of diverse immune function categories described in panel A, using canonical correlation analysis. Correlations are shown separately for each immune gene category. Corresponding statistics, including canonical R-square and R-squares representing only significant pathogen-specific trends, are also shown for each immune gene category.

To further explore these associations, we performed a canonical discriminant analysis to obtain a linear combination of expression values of immune-related genes, separating the effects of infection across pathogen types and selection regimes in each immune category. We corroborated the statistical differences due to infection treatment as found in MANOVA. However, the effects were pathogen-specific, highlighting the mechanistic differences in host responses across pathogens. For instance, while infection produced comparable patterns of changes in gene expression values in both P- and M-beetles across all the immune categories, B-beetles showed contrasting patterns of changes in at least two of these functional categories, namely inducible immune effectors and immune receptors (note the opposite direction of changes with Bt vs Pe and Mx infections; **Fig 5B; Tables S14**–**18**). These immune gene expression patterns plausibly reemphasize the greater functional overlaps in immune repertoires between M- and P-beetles relative to B-beetles (**Fig. 5B; Tables S14**–**18**). We also found significant interactions between the selection regime and infection treatment in some functional immune categories where infection affected evolved beetles differently from their respective control populations (**Fig. 5B; Tables S14**–**18**). For example, Pe infection induced more divergence in the expression patterns of inducible effectors, receptors, and immune regulators in evolved P relative to C beetles (e.g., note the divergence between sham-infection vs bacterial infection; **Fig 5B, Tables S14**–**18**). It also induced changes in phenoloxidase response-related genes that were initially non-responsive in control beetles. In contrast, evolution against Bt produced changes limited to only fast-acting melanisation response and ROS. Interestingly, evolution against Mx infection involved changes in inducible immune effectors and phenoloxidase responses, suggesting the involvement of molecules partially involved in both P- and B-regimes, although their functional implications might differ.

Finally, we applied canonical correlation analyses, followed by linear regression analyses, to determine whether the observed changes in the gene expression profile of the aforementioned immune categories predicted the phenotypic variations between the control vs selected regimes across pathogens. In each case, we used a joint estimate of the bacterial load of individual beetle hosts and the infection susceptibility, estimated as the hazard ratios (*36*) of infected vs sham-infected beetles (where Hazard ratios greater than 1 denote higher mortality in the infected beetle), to gain an integrated view of post-infection fitness outcomes as a function of pathogen burden and concomitant survival costs. We assumed significant correlations between gene expression values and the phenotypic changes to imply whether the concerned category of immune molecules can explain the observed variation in phenotypic traits during experimental evolution. Overall, phenotypic variations of Pe-infected beetles correlated with a maximum number of immune categories, including receptors, immune regulators, and inducible immune effectors, followed by Bt-infected beetles that correlated with regulators and melanisation responses, and Mx-infected beetles that correlated with only receptors (**Fig. 5C; Tables S19**–**23**). Similar patterns also emerged when we separately analysed the associations of infection susceptibility with the gene expression profile using linear regression analyses. Pe-infected beetles still had more correlations (i.e., with both receptors as well as melanisation response) relative to B and M beetles that either correlated with melanisation response or receptors, respectively (**Fig.S9A; Table S24**). In contrast, no such correlations existed with the bacterial load of Bt-infected beetles, as opposed to Pe- and Mx-infected beetles, where, in addition to exhibiting correlations to receptors and regulators respectively, they also showed association with melanisation response (**Fig. S9B; Table S25**). While these correlations suggest a larger scope of the modulating immune responses at various functional levels in P-beetles to increase post-infection fitness, they possibly also reemphasized the potential divergence of immune strategies adopted in B-beetles from that of P- and M-beetles to control the pathogen growth.

In addition, we also used KEGG enrichment analyses to reveal broad similarities in how several key metabolic pathways responded against Pe and Mx infection (**Fig. S10**). For instance, unselected C-beetles infected with Mx and Pe showed downregulation of glutathione and several components of amino acid (e.g., valine, leucine, and isoleucine), and carbohydrate (e.g., amino sugar and nucleotide sugar) metabolism. They also showed downregulation of glycolysis and upregulation of phagosome maturation pathways, which contrasted with Bt-infection. In the evolved M- and P-beetles, we noted a downregulation in both the citrate cycle and OXPHOS pathway (**Fig. S10**). Also, while evolved B-beetles overexpressed mismatch and nucleotide excision repair pathways, both M- and P-beetles produced no changes in their expression. These results thus suggest the possibility of common metabolic bases underlying overlapping immune responses against Mx and Pe.

## DISCUSSION

Despite the ubiquity of coinfections and their direct relevance to many infectious diseases (*2*, *4*), it is unclear how they influence host adaptive trajectories against pathogens and concomitant immune system evolution. Also, what are the specific drivers of such evolutionary effects of coinfections? Here, we used mathematical models to propose within-host growth rate, the rate of virulence manifestation, and the infection-driven mortality window of individual pathogens (i.e., between first and last host mortality) as critical determinants of adaptive success against coinfections. In the absence of strong competitive interference or cross-reactive immunity between pathogens (*27*, *28*), rapidly growing pathogen counterparts, imposing acute mortality surges early in the infection, determined the course of evolution against coinfections. Moreover, if such an early surge of survival costs is also expressed rapidly within a short infection window, the rate of adaptation against coinfections might be delayed. We hypothesize this as a possibility that arises when appropriate immune responses are unavailable or cannot be induced against such fast-acting pathogens within the available infection window to curb their acute early-infection costs (*31*, *37*). Subsequently, we empirically validated whether fast-growing pathogens can drive the adaptation against coinfection, using replicated populations of *T. castaneum* evolving against bacterial pathogens of distinct Gram-types (i.e., Bt and Pe) with contrasting within-host dynamics, virulence manifestation rates and differential host immune modulations (*38*, *39*). Bt grew faster early in the infection, inducing early and rapid mortality surge within 12h (i.e., fast-acting), followed by rapid clearance by the host. In contrast, Pe grew relatively slowly, causing long-lasting persistent infections with mortality beginning around 24h post-infection (i.e., slow-acting). We found the rate of adaptation to be fastest against Pe, with half of the replicate populations evolving resistance as early as generation 8, whereas resistance evolution against fast-growing Bt was delayed the most. Also, as predicted by the model, the rate of adaptation against coinfection by Mx indeed appeared to be constrained by fast-acting Bt such that M-beetles followed almost a similar evolutionary trajectory as that of beetles infected with Bt alone (e.g., most replicate populations taking 15–22 generations to evolve resistance). Another striking aspect is that while the survival success of the P-beetles rose to ∼90%, the survival of both M- and B-beetles could not increase beyond ∼75% despite a continuous strong selection for 30 generations. This possibly indicates the constraints associated with evolving resistant alleles against Bt cells that are present in both M- and B-beetles during their early infection phase, restricting their net fitness gain to much below that of their P-beetle counterparts (*40*, *41*).

Here, an emerging question is— how might Bt cells drive the dynamics of adaptive evolution against Mx? We noted that beetles infected with Mx showed a sharp decline in survival early in infection (between 16–20h), which broadly resembles the mortality pattern of beetles that were only infected with Bt. Also, beetles that succumbed to infection within this early timeframe predominantly carried many Bt cells (∼10^5^–10^6^ cells/female), linking the overgrowth of Bt to lethal infections. Interestingly, the estimated levels of the bacterial load causing such terminal infection did not correlate with the time post-infection at which death occurs. Hence, they also denoted the maximal Bt load that beetles could tolerate before they died (*20*). Several dead beetles also carried Pe cells, but neither their frequency nor their within-host Pe density was sufficient to explain all the beetle mortality observed during the early infection phase, hinting at the limited role of Pe in driving the early survival costs of coinfection. In contrast to dead beetles, surviving beetles early in infection either had a much lower Bt burden than their dead counterparts or cleared the infection below the detection level within 20h. The ability to restrict the growth of Bt below their threshold density, which otherwise could lead to terminal infections, followed by rapid clearance, was thus critical for these beetles to survive the early phase of coinfection. These results also conform to recent studies with *D. melanogaster,* where similar binary infection outcomes have been reported across pathogens (*20*, *21*, *26*), underscoring the pivotal roles of rapidly induced immunity in effectively curtailing the pathogen overgrowth early in infection.

As expected, the ability to prevent Bt overgrowth early in coinfection also increased in beetles adapted against Mx infections. When challenged with Mx-infection, fewer individuals from evolved M-populations carried the lethally high Bt density (∼10^5^–10^6^ cells/ beetle) relative to C-beetles, thereby explaining the reduction in their early-infection mortality. However, increased survival of evolved beetles was not achieved by merely clearing the Bt cells, as their number, by and large, did not vary considerably between the live M- vs C-beetles. Subsequently, C-beetles that survived the infection could clear the Bt cells at a nearly equal rate to that of M-beetles. This suggests that pathogen selection did not improve pathogen clearance ability in evolved beetles. Instead, it could have favoured mechanisms to arrest the Bt growth below the critical density, otherwise leading to lethal infections (*20*). This is likely also the reason why transcriptome analyses of beetles challenged with only Bt infection had several differentially expressed immune effectors upon infection (e.g., AMPs Attacin 2, Coleoptericin and Tenecin 1) (*42*) and still, none of them responded differently in evolved B beetles, suggesting no added contribution towards experimental evolution against Bt. Also, the overall changes in the gene expression profile of different immune effector groups, including AMPs, PO or ROS, did not explain the variation in the overall bacterial load across beetle lines. Perhaps more relevant changes in Bt-resistant beetles were detected in terms of their higher basal expression levels (i.e., without infection) of apolipophorins, possibly facilitating phagocytosis and pathogen pattern recognition (*43*) or chymotrypsin, which is known to arrest the growth of Gram-positive bacteria (*44*), including neutralization of Bt-toxins (*45*). Increased circulation of these molecules, even in the non-immune challenged state of B beetles, might thus play a more important role in early detection and prevention of Bt overgrowth.

By contrast, immune strategies against Bt in M-beetles may be more complex due to confounding effects of immune responses against chronic Pe infections persisting throughout the oviposition window of these experimentally evolving beetles (i.e., 3–8 days post-infection). Moreover, unlike Bt infection, surviving M-beetles consistently had reduced Pe load relative to C-beetles, suggesting the potential immune activation against Pe to minimize the infection costs while reproducing (*46*).

Finally, despite receiving a lower infection dose (M- vs P-beetles: ∼10^3^ vs 10^4^ cells/female), Pe cells in M beetles grew at an equivalent level as that of P beetles (∼10^5^ cells/female within 12 hours), which indicates that both beetle populations might have eventually experienced similar selection pressure from the severity of Pe infection. These hypotheses were further corroborated by comparing the reproductive costs of each infection type in the unselected C-beetles (see **Fig. S11, Table S26**). In the case of both Pe and Mx infections, the persistence of Pe cells during the oviposition window was also associated with a reduction in reproductive outputs. However, this contrasted with beetles challenged with Bt infection. Bt-infected beetles that survived until the oviposition window reproduced as much as their uninfected control counterparts, possibly attributed to their ability to clear the infection completely by then. Based on these observations, we thus speculated strong selection pressure on both M- and P-beetles from the beginning of their selection treatment to evolve counterstrategies to reduce the reproductive costs imposed by a common pathogen that persists longer inside the host (*47*). More specifically, to this end, M-beetles might evolve more similarities with P-beetles vis-à-vis their immune responses rather than temporally compartmentalizing immunity against individual participating pathogens (*48*). Our transcriptome analyses that revealed larger overlaps in the set of genes and their expression profile against Pe and Mx infection, both before and after experimental evolution, perhaps supported this idea.

The possibility of mechanistic congruence between M- and P-beetles is further highlighted by the linear discriminant analyses of immunity-related gene expression data (*49*). In fact, many of them, classified into various functional categories ranging from sensing the pathogen or pathogen-associated molecular patterns (receptors) and regulating the immune responses to immune effectors such as AMPs, lysozyme, phenoloxidase cascade and ROS production (*34*, *50*), showed more concurrent gene expression patterns between M- and P-beetles. These patterns, however, did not always match with that of B beetles, as some of these functional categories, such as AMPs and lysozymes, immune regulators and receptors, showed changes either in the opposite direction to that of M- or P-beetles or produced no changes after infection. Also, unlike in M- and P-beetles, none of the immune groups correlated with the changes in the overall bacterial burden before and after the evolution against Bt. Together, all these patterns thus hint at distinct functional implications of these immune groups in B-beetles relative to both M- or P-beetles.

The similarity in immune responses against Pe and Mx infection was also reflected by their resemblance in metabolic changes. For example, KEGG enrichment analyses revealed the downregulation of several important components of carbohydrate (e.g., glycolysis, amino sugar and nucleotide sugar metabolism) and amino acid (e.g., valine, leucine and isoleucine) metabolism in unselected C beetles (*42*). Besides, evolved P- and M-beetles showed reduced OXPHOS metabolism and increased glycolytic enzyme hexokinase 2 expression, suggesting shifting energy metabolism to support immune activation in these beetles (*51*, *52*). However, such metabolic patterns were reversed in B beetles, which could corroborate why we failed to detect increased expression of immune effectors after experimental evolution. Instead, the enrichment of pathways related to increased DNA repair (*53*) and phagosome maturation (*54*) might indicate strategies to reduce the DNA damage caused by immune activation (*55*) and the use of alternative immune strategies in B beetles (e.g., cellular immunity (*38*, *56*)) respectively.

Finally, a detailed comparison of phenotype-by-immune gene expression correlations across pathogens and infection types offered critical molecular insights into their divergent adaptive dynamics. For example, strong correlations between the combined phenotypic changes (i.e., post-infection survival and bacterial load) in P-beetles and diverse categories of immune-related molecules such as receptors (e.g., PGRP SC1a/b-like, PGRP2 (*33*)), regulators (e.g., Relish (*57*)) and inducible effectors (including Attacin 1, Attacin 2, Tenecin 1 and Ctenidin 1 (*34*)) might underscore a wider scope for selection, acting parallelly and more effectively across various functional levels of their immune signalling cascade (*58*, *59*). This, in turn, can accelerate their rate of adaptation. This notion can also be supported by previous analyses where immune molecules, particularly those involved in pathogen recognition and immune regulation, have been shown to evolve more rapidly under strong positive selection than other non-immune genes (*60*). Moreover, the multi-level immune crosstalk between receptors, regulators and effectors driving phenotypic variations against Pe corroborates the assumptions of our theoretical model. For instance, faster adaptation against slow-acting Pe-like pathogens was possible because slower mortality costs expressed over a prolonged infection window enabled beetles to employ functionally more diverse phenotype-by-immunological modulations under pathogen selection (*25*, *61*).

In contrast, the scope for such phenotype-by-immunological modulations in M- and B-beetles was limited. For example, unlike P-beetles, their immuno-competence phenotype correlated with either receptors or regulators but not with both, which might reduce the number of potential loci available to evolve rapidly under selection. Scopes for selection might be even more constricted in B beetles, as their phenotypic variations correlated with PO response (*62*), which, in addition to serving as a critical insect immune defence component, exerts multiple pleiotropic roles in insect physiology (*63*–*65*). For example, the observed correlation was mainly driven by reduced PO enzymatic activity in evolved B-beetles (**See SI, Fig. S12; Table S27**), conforming with their lower expression levels of phenoloxidase 2 and tyrosine decarboxylase transcripts (*63*, *66*), but evolution via such decline in PO activity might also impose development and reproductive costs (*63*, *65*). Besides, evolved B beetles also showed more divergent expression profiles of ROS-related genes after Bt infection than the unselected beetles, driven primarily by down-regulation of Glutathione S-transferase 1 after Bt infection, which might incur higher cytotoxicity (*67*), thereby adding significant costs to fast adaptation against Bt.

In summary, this study present a unique integrated framework, combining theory and experiments, to identify drivers of host adaptive dynamics against coinfecting pathogens. Note that we could test only a few specific infection conditions amidst numerous other possible interactions between host and coinfecting pathogen types and their diverse infection outcomes. Yet, the coherence between theoretical predictions and our empirical datasets, while establishing the role of pathogen growth dynamics and virulence manifestation patterns in driving phenotypic and mechanistic trajectories, indicated the broader implications of our findings. Another striking outcome of our work is the decoupling of the overall rate of phenotypic evolution vs mechanistic bases against coinfecting pathogens relative to the effects of individual pathogens. This eventually highlighted the asymmetry in why and how individual pathogens might unequally bias the adaptive dynamics against coinfection vs underlying genetic mechanisms rather than their simple additive effects (*68*), offering exciting avenues for future theoretical models to encompass other infection types and more mechanistic explorations. Finally, our systematic investigation of host adaptations against multiple pathogens and infection contexts in a single comparative framework may instigate more fundamental work to fill the gaps in our understanding of how innate immune features might evolve across pathogens and infection types.

## MATERIALS AND METHODS

### Mathematical simulation of host survival and adaptation against coinfecting pathogens with contrasting growth and virulence dynamics

To model the effects of coinfections caused by two pathogens with contrasting growth dynamics on host survival and adaptive responses, we began by simulating their growth dynamics following a theoretical framework described previously by Duneau et al. (*20*) (Detailed descriptions of parameters used in these simulations, as well as those described below, are provided in supplementary methods). The model, originally built upon experimental data from fruit flies, assumed uninhibited pathogen growth initially followed by either host immune response clearing the pathogen or the host succumbing to acute infection, leading to binary outcomes — a phenomenon validated empirically in other species as well, including mice and flour beetles (*69*, *70*). To capture similar growth dynamics, we thus combined two demographic models where we first simulated the divergent pathogen growth patterns (i.e., rapid vs slow) without the interference of host immunity, based on the Baranyi model (*71*). We then used an exponential decrease model to simulate the pathogen clearance where host immunity could either clear the infection completely or maintain a lower pathogen burden, producing long-lasting infections (*26*). Note that we did not consider the pathogens that cause lethal infections causing complete mortality or benign infections, as both conditions might preclude the possibility of host adaptations against them. Hence, we could simulate pathogens with the following 4 distinct growth dynamics: e.g., rapidly growing pathogens causing acute infections, followed by (a) rapid clearance (Rc) or (b) persistent infection (Rp); Slow-growing pathogens causing acute infection, followed by (c) rapid clearance (Sc) or (d) persistent infection (Sp). Subsequently, we paired pathogens with only contrasting growth rates (rapid vs slow) to simulate the following coinfection scenarios: e.g., Rc-Sc, Rc-Sp, Rp-Sc, and Rp-Sp.

Here, we note that coinfecting pathogens might compete for limited resources (*27*), inhibit each other’s growth by producing toxins (*12*) or induce cross-reactive immune mechanisms (*28*), which can influence their within-host growth dynamics. To model such effects, we thus described a coefficient of interference *α*

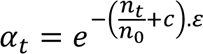

Here, we assumed α to range from 0 to 1, denoting the maximum or no interference between the coinfecting pathogens respectively. Also, since interference due to competitive inhibition and immune cross-reactivity might proportionally increase with the load of the interfering pathogen, the magnitude of *α* at time *t* is inversely proportional to the ratio of load of the interfering pathogen at time *t* (*n_t_*) and the initial bacterial load (*n_0_*). Additionally, we note the possibility that residual interference may persist even after the interfering pathogens are cleared by the host immunity. To account for this effect, we introduced an offset factor *c*, denoted as the ratio of initial bacterial load (*n_0_*) and maximum bacterial load (*n_max_*) of the interfering pathogen. Finally, we also described a coefficient *ε*, combining the net interference posed by increasing resource-driven competition or toxin-mediated inhibition between pathogens (*β*) and/or immune cross-reactivity (*γ*) mediated via activation of common host immune components across pathogens, that is inversely proportional to the value of *α* (such that *ε* = 0 or 1 represents no interference or maximum interference respectively; see SI for details).

Next, we predicted the post-infection survival probability of hosts challenged with a combination of rapid (R)- vs slow (S)-growing pathogens based on their simulated within-host growth dynamics and the estimated interference coefficient as described above:

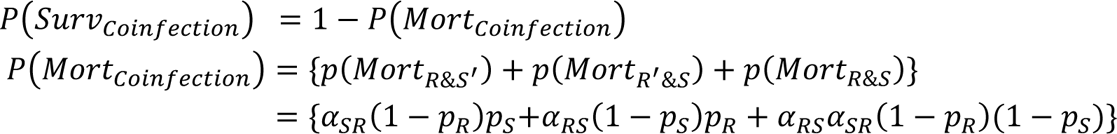

Here, we first conceptualized the host survival probabilities under coinfection *P(Surv_coinfection_)* as individuals that could avoid the mortality induced by individual pathogens (Mortality probabilities: *p*(*Mort*_*R*&*S*′_) or *p*(*Mort*_*R*′&*S*_) against R vs S pathogens) as well as their combined actions (*p*(*Mort*_*R*&*S*_)). Here, *S’* and *R’* denote complementary terms of probability. We then transformed these mortality probabilities as a function of survival probabilities against individual pathogens (i.e., *p_R_* = survival probability against R; *p_S_*= survival probability against S) and their coefficient of interference (*α)*. We assumed that *α* is directional so that *α_SR_* describes the effect of ‘S’ on ‘R’ and vice-versa (*α_RS_*).

### Modelling the rate of adaptation

We note that the survival costs against coinfections can increase with reductions in competitive interference or immune cross-reactivity between pathogens (i.e., high *α*; low *ε*). Moreover, they follow the predicted survival patterns of the rapidly growing pathogen counterpart during the early infection phase. We thus chose to model the combinations of coinfecting pathogens only with low interference, which is likely to posit strong selection pressure on the hosts to first counter the early infection costs of rapidly growing pathogens. We also assumed that the ability to deploy and evolvability of effective immune responses could be constrained by the response time available to the host after infection (*21*, *30*). For example, rapidly proliferating pathogens causing acute-phase infection and a rapid surge of mortality manifested within a shorter timeframe after infection (e.g., Rc) might outrun the host immunocompetence due to limited immune repertoire availability or failure to induce or replenish appropriate levels of required immune responses. Consequently, host adaptative success against such pathogens can be constrained. In contrast, hosts infected with slow-growing pathogens that have prolonged infection windows with sustained survival costs (e.g., Sp) can also afford a longer response time to deploy and modulate various immune components (e.g., both constitutive vs inducible responses (36)) against pathogens. We thus assumed that the ability to respond and modulate effective immune responses to counter the infection could be directly proportional to the total infection window, where the pathogens first proliferate to cause the acute infection phase, followed by the host mortality window (i.e., the time between the first and last post-infection mortality). We modelled the efficacy of host adaptations against individual pathogens as a function of the relative scope for immune modulations within their total effective infection window, expressed as a ratio with respect to the total generation time of the host (e.g., conceived as early development to time to reproduction to initiate the next generation).

Since rapid-growing pathogens imposing early virulence manifestations drive the outcome of coinfection during the early infection phase, we next assumed that the time taken to inflict the first mortality after infection could be comparable between the coinfection vs rapidly growing pathogens. Consequently, we modelled the efficacy of adaptation against coinfections as a function of the available immune modulations only during the mortality window of rapid-growing pathogens with respect to the total mortality window caused by both pathogens. In all these cases, we expected that the host adaptive trajectory would initially have a lag period where survival advantage cannot be detected, followed by an increase until it reaches an asymptote, with no further gain in survival advantage. We thus used a logistic model to predict the adaptative trajectories, formulated as:

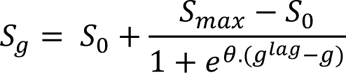

Here, we describe the survival against infections caused by single pathogens or coinfections at generation *g* as *S_g_*, which depends on (a) the survival of the ancestral populations (*S_0_*) and their maximum gain in survival (*S_max_*) after successful adaptation against pathogenic infections; (b) the minimum number of generations elapsed (*g^lag^*) before hosts could show survival advantage; and (c) the parameter *θ*, denoting the constraints of evolving appropriate immune responses within the host mortality window for single or coinfecting pathogens (detailed in SI methods). Overall, the parameters *θ* and *g^lag^* influence the variable trajectories of host adaptation against single vs coinfecting pathogens. We simulated all the parameters using R.

### Experimental quantification of virulence and growth dynamics of coinfecting pathogens

We used a large, outbred population of *Tribolium castaneum* (*72*) adapted to laboratory conditions for >2 years before commencing the experiments (see SI for more details on baseline population maintenance and assays described below). Also, based on the observations from other experiments (*32*, *73*), we chose two naturally relevant bacterial entomopathogens that are likely to show contrasting growth dynamics and rates of virulence manifestations within insect hosts: fast-growing *B. thuringiensis* DSM2046 (Bt) (*74*) vs slow-growing *P. entomophila* L48 (Pe) (*75*), causing a rapid vs slower onset of mortality respectively. To quantify their effects on beetle hosts, we pricked 10-day-old virgin females from baseline populations with a needle dipped in a bacterial slurry comprised of either Bt (∼ 8×10^^7^ cells/µl) or Pe (∼ 4×10^^9^ cells/µl), or a mix (1:1) of both bacterial cells (Mx) and monitored their survival for 10 days (See SI for input infection dose). We used sham-infected beetles pricked with sterile Insect Ringer solution (*74*) as procedural control for our infection assays (n= 30 females/infection treatment). We also tracked the changes in growth dynamics of these pathogens inside surviving beetles sampled at regular intervals for the next 24h for Bt (or ∼3 days for Pe), both in the context of infections caused as individual vs co-occurring pathogens (n=8–10 replicates with pooled homogenate of 3 females/infection treatment/time point), using established protocols in the lab (*76*). We differentiated the Bt and Pe cells by their distinct colony sizes and morphologies on the Luria agar plates (See SI methods). For each pathogen, we analysed the (a) post-infection survival data using Cox proportional hazard analysis (*77*), using infection treatment as a fixed effect in “survival” package in R (*78*), and (b) log-transformed bacterial load dynamics data using a generalised linear model (‘glm’ function in R) fitted to a gamma distribution with infection treatment and time of bacterial load estimation as fixed effects. Separately, we also assayed the bacterial load of a subset of females that succumbed to Mx-infection within the first 48h to estimate the bacterial load upon death and the relative contribution of Bt vs Pe burden in causing mortality during coinfection (n= 34 females).

### Experimental evolution paradigm

Next, we used the experimental evolution paradigm for 30 successive generations to examine the adaptive dynamics against coinfecting pathogens (*32*). We used the baseline beetle population to create five selection regimes: namely (I) Unhandled regime (U-regime): populations that did not undergo any treatment; (II) Control regime (C-regime): Unselected control populations sham-infected with sterile Ringer; (III) Infected with Bt (B-regime); (IV) Infected with Pe (P-regime); (V) Infected with a mixed culture of both Bt and Pe (M-regime), with each of these regimes having four independently evolving replicate populations (i.e., C1–4, B1–4, P1–4 & M1–4; See SI for infection doses). We infected (or sham-infected) 9–10 days old virgin adult male and female beetles (*32*).

Three days later, we combined the surviving beetles into 75 pairs and allowed them to oviposit for another 5 days (i.e., beetle reproductive window; day 3–8 post-infection). Note that although we maintained ∼75 breeding pairs for each selection regime, we infected an excess of virgin beetles (∼300–400 beetles/ replicate population) for every generation, as we expected high mortality after the respective infection treatments. During experimental evolution, this ensured we had sufficient individuals to set up the 75 mating pairs to oviposit every generation. We adjusted our infection doses to induce ∼60–65% mortality within 8 days post-infection across infection treatments, thereby enabling us to initiate beetle lines with comparable selection pressure across pathogen-selected regimes. After three weeks of egg incubation, we isolated male and female pupae from each population and allowed the eclosed adults to initiate the next generation after the relevant selection treatment. We handled the four replicate populations from each selection regime on different days (but C_i_, B_i_, P_i_ and M_i,_ where i=1–4, were handled together on the same day) and maintained continuous divergent pathogen selection. For each of the 4 replicate populations across selection regimes, we also estimated the proportion of the surviving adults (out of the total number of infected beetles) pre- (i.e., day 3 post-infection) and post-reproductive window (i.e., day 8 post-infection) at every generation (except the first two generations) to track the overall changes in survival post-infection across selection regimes. Moreover, to understand the contrast between diverging evolutionary trends across generations from different selection regimes, we used a generalised linear model fitted to Gaussian distribution followed by performing pairwise comparisons using “emmeans”.

### Quantifying the evolved responses against coinfection

To understand the dynamics of adaptation against single pathogens vs coinfections, we repeatedly assayed post-infection survival of each replicate population across selection regimes against their respective infection treatments during experimental evolution and compared with that of control unselected beetles (i.e., C vs P; C vs B or C vs M beetles after Pe, Bt and Mx infection respectively), at multiple generations (e.g., generations 8, 13, 15, 18, 20, 22 and 28). We used 9–10 days old standardised females (for logistical reasons, we could not test males, except generation 28 when both sexes were assayed) (n=24–60 beetles/regime/population/generation), collected after one generation of relaxation of pathogen selection from the generation of interest, to minimise the transgenerational effects (*32*). For each pathogen-selected regime, we compared them separately with the C-regime after the respective infection treatments, using the mixed-effects Cox model (with the selection regime as a fixed effect and replicate populations as a random effect) (*77*), followed by analysing each replicate population separately, using Cox proportional hazard analyses (with the selection regime as a fixed effect). Besides, we also quantified the bacterial load of evolved beetles when all the replicate populations showed improved post-infection survival (P- and M-beetles at generation 18; B-beetles at generation 22) to test whether their improved survival can be explained by lower bacterial burden relative to their control counterparts (N= 10–15 replicates with pooled homogenate of 3 females/selection regime). Since different pathogens might manifest their virulence at different rates with divergent within-host pathogen growth dynamics, we sampled 9–10 days-old females from B and P regimes (and their corresponding Control regimes) after the onset of the first 10–15% mortality (information derived from post-infection survival curves) after respective infection treatments. For the M regime (and its corresponding control), we sampled females at two time points to obtain an adequate number of both Bt and Pe cells (See SI for detailed methods and analyses).

However, to explain the mortality patterns in more detail, we next (at generations 26–28) characterized the dynamics of within-host bacterial load in one of the replicate populations from both evolved and control beetles by assaying 9–10 days-old standardised females every few hours (See SI for detailed methods). We tracked Bt cells in B vs C regimes until 18 hours (or Pe cells in P vs C regime until 72 hours), whereas, for Mx infection, both the bacterial cells were assayed until 50h (n= 6–10 replicates with pooled homogenate of 3 females/selection regime/time point/infection treatment). Simultaneously, we also noted the beetle death during these experiments and estimated their bacterial load as soon as they succumbed to respective infection treatments across selection regimes to understand the link between growth dynamics, virulence manifestation, and the maximum pathogen load that beetles could tolerate before death (n=13–80 replicates/selection regime/infection treatment). We analysed the bacterial load data using a generalised linear model fitted to a gamma distribution, with selection regime and time of assay as fixed effects for live beetles and only selection regime as fixed effects for dead beetles.

### Transcriptome analyses

Finally, to gain mechanistic insights into the evolved responses of P-, B- and M-beetles, we performed whole-body transcriptome analyses of 10-day-old virgin females after the respective infection treatments (or sham treatment) (n= 4 replicates; each comprised of 10 females pooled together from each replicate population/infection treatment/selection regime). Here, we sampled females from each infected population and their sham-infected counterparts across the control and three pathogen-selected regimes (See SI methods for more details on experimental protocol and analyses). Attributed to divergent temporal growth and virulence dynamics across pathogens and infection types, we extracted the RNA from beetles after the onset of the virulence manifestation (i.e., 10–15% mortality around 24h, 8h, or 16h after Pe, Bt and Mx infection respectively) to obtain gene expression profiles at comparable fitness variations, rather than at a specific time point post-infection. We used Qiagen RNeasy Minikit to extract RNA following the manufacturer’s protocol. We sent the isolated RNA samples to a commercial sequencing facility (Neuberg Diagnostics Private Limited) for downstream processing. The quantity and quality of extracted RNA was checked on a Qubit 4.0 fluorometer (Thermofisher #Q33238) using an HS RNA assay kit (Thermofisher #Q32851) and on TapeStation using HS RNA ScreenTape (Agilent), respectively. The libraries was prepared using TruSeq® Stranded Total RNA kit (Illumina #15032618, Illumina #20020596) post poly-A enrichment (*79*). The sequencing was performed on an Illumina Novaseq 6000 platform using a 150 bp paired-end chemistry. We analyzed the transcriptome data using an HTseq-based customized pipeline (*80*) and Tcas5.2 as the reference genome (*81*). We estimated differential gene expression (DEGs) using R package “DESeq2” (*82*). We used “pheatmap” and “RColorBrewer” packages in R to visualize expression profiles of all the DEGs by a pooled-population heatmap based on the z-score of normalized read counts. We counted the common vs unique sets of up- and down-regulated DEGs, including those with contrasting variations, among the three pathogen-selected regimes and visualized them using R-package “ComplexUpset”. We performed a principal component analysis based on the normalized count data of all the DEGs from all three pathogen-selected regimes. Subsequently, we obtained all the gene ontology terms (GO terms) using Blast2Go. GO terms for KEGG were used to perform pathway enrichment using the R-package “gProfiler2” and visualised using the R-package “ggplot2”.

Subsequently, to understand the possible role of immune responses, we categorised DEGs with known immunological roles into five broad functional categories: (a) pathogen and immune receptors; (b) immune regulators; (c) inducible immune effectors, including AMPs and lysozymes; fast-acting constitutively expressed (d) melanisation response involving phenoloxidase pathway; and (e) production of reactive oxygen species. We estimated a linear combination of gene expression profiles for each immune category as a function of infection treatments and selection regimes, using canonical discriminant analysis (*49*, *83*). Further, to understand how these gene expression profiles correlated with phenotypic variations (either individually with the hazard ratios (as a proxy of survival response) or bacterial load and as a combined estimate of both hazard ratio and bacterial load for each infection treatment and selection regime), we performed linear regressions and canonical correlation analysis (*49*, *83*) respectively.

## Conflict of interest

We have no conflict of interest.

## Author’s contributions

Conceptualisation: Imroze Khan, Srijan Seal, Dipendra Nath Basu

Design of the experiment: Imroze Khan, Srijan Seal, Dipendra Nath Basu, Ishaan Gupta, Triveni Shelke

Data curation: Srijan Seal, Dipendra Nath Basu

Formal analysis: Dipendra Nath Basu, Srijan Seal, Rintu Kutum

Funding acquisition: Imroze Khan, Srijan Seal, Ishaan Gupta

Investigation: Srijan Seal, Dipendra Nath Basu, Kripanjali Ghosh, Aryan Ramachandran, Imroze Khan

Supervision: Imroze Khan

Visualisation: Dipendra Nath Basu, Srijan Seal,

Writing – original draft: Imroze Khan, Dipendra Nath Basu, Srijan Seal

Writing – review & editing: Imroze Khan, Dipendra Nath Basu, Srijan Seal, Rintu Kutum

## ACKNOWLEDGEMENTS

We acknowledge Deepa Agashe, Shivani Krishna, Sandeep Ameta, Basabi Bagchi, Saubhik Sarkar, Biswajit Shit, Shriya Palchaudhuri and Nilabhra Mitra for their feedback on the manuscript. We thank Gautam Menon and Abhirup Banerjee for providing their critical insights during mathematical modelling.

## FUNDING SOURCES

We thank the DBT-Wellcome Trust Intermediate Fellowship (IA/I/20/1/504930 to IK), SERB-DST (ECR/2017/003370 to IK), Society for Study of Evolution (R.C. Lewontin’s Early career research grant to SS), IITD-Ashoka MFRIP Funding (MI02416G to IK and IG) and Trivedi School of Biosciences at Ashoka University (Simon-Ashoka early career fellowship to DNB) for funding this research.

## Notes

### Competing Interest Statement

The authors have declared no competing interest.

