## Supplementary materials for "PATHOGEN GROWTH AND VIRULENCE DYNAMICS DRIVE THE HOST EVOLUTION AGAINST COINFECTIONS"

### SUPPLEMENTARY METHODS

### A. Modelling host survival and adaptation against coinfections

Simulating pathogen growth dynamics

We started by simulating the growth dynamics of pathogens following a theoretical framework described by Duneau et al. 2017 (*1*). To simulate diverse within-host growth dynamics and pathogen clearance, we first described the pathogen growth patterns (i.e., rapid vs slow) without the interference of host immunity, based on the Baranyi model (*2*) with the following parameters: (a) initial bacterial load ($n_0$); (b) the maximum bacterial load sustained by a host ($n_{max}$); (c) the maximum growth rate ($\mu$); (d) and the lag time to growth ($t^{lag}$). Upon obtaining the mean of the estimated bacterial load across different time points from the model, we used a standard deviation ($\sigma_b$) to simulate the pathogen load from individual hosts, following a log-normal distribution. Subsequently, we also described a time window after infection ($t_c$) when host immunity is activated to prevent pathogen growth and a probabilistic value ($p_s$) of the binary outcomes where bacterial growth either continues to follow the Baranyi model or deviates from its predicted trajectory. Further, we randomized these deviating individuals and simulated their pathogen clearance using the exponential decrease model based on the pathogen load ($n_c$) at $t_c$ and the rate of pathogen clearance thereafter ($\delta$), such that the pathogen gets either entirely cleared by the host or persists in the host body at a lower pathogen burden (incomplete clearance), resulting in long-lasting infections (*3*). We simulated pathogens with the following 4 distinct growth and clearance dynamics: e.g., rapidly growing pathogens causing acute infections, followed by (a) rapid clearance (Rc) or (b) persistent infection (Rp); Slow-growing pathogens causing acute infection, followed by (c) rapid clearance (Sc) or (d) persistent infection (Sp). Note that we did not consider pathogens causing complete mortality or benign infections as, in both cases, selection may not act effectively for host adaptation. To obtain the effects of coinfections, we paired the pathogen types with only contrasting growth rates (rapid vs slow): e.g., Rc-Sc, Rc-Sp, Rp-Sc, and Rp-Sp.

Parameterising the mode of interference (ε)

We note that the net interference between pathogens can increase (i.e., lower $\alpha$-values) with higher resource-driven competition or toxin-mediated inhibition between pathogens (collectively represented as $\beta$) and/ or immune cross-reactivity mediated via activation of common host immune components across pathogens (characterised by a higher ratio of common vs unique immune components activated by the interfering pathogen against the other pathogen counterpart; represented as $\gamma$). We conceived the value of $\alpha$ as inversely proportional to both $\beta$ and $\gamma$, combined into a ratio-metric parameter ε described as:

$$\varepsilon = \frac{(\beta + \gamma)}{k}$$

We represented $\varepsilon$ as a scaled summation of $\beta$ and $\gamma$, where $k$ is the scaling factor, ranging between 0 and 1 (while $\varepsilon$ = 0 represents no interference; $\varepsilon$ = 1 denotes maximum interference between pathogens from competitive inhibition and immune cross-reactivity).

Parameterising the simulations of host adaptative trajectories against individual vs coinfecting pathogens

As described in the main text, to model the host adaptation against both single- and coinfecting pathogens, we parameterized the following logistic model representing survival advantage in each generation, formulated as:

$$S_g = S_0 + \frac{S_{max} - S_0}{1 + e^{\theta.(g^{lag} - g)}}$$

Where the survival against individual pathogens at generation $g$ is denoted as $S_g$, which depends on (a) the survival of the ancestral populations ($S_0$) and their maximum possible survival benefit ($S_{max}$) after successful adaptation against the pathogens; (b) the minimum generations ($g^{lag}$) required before hosts exhibit a survival advantage against pathogens; and (c) the parameter $\vartheta$, denoting the constraints of evolving appropriate immune responses within the host mortality window.

However, we described the parameters $g^{lag}$ and $\theta$ separately for infections caused by individual vs coinfecting pathogens.

(a) Adaptation against individual pathogens:

$$\theta = f.\zeta.G_t^{-1}$$

$$g^{lag} = (l.\psi^{-1} + m.\zeta^{-1}).G_t^{-1}$$

Here, we assumed that the host adaptation rate against individual pathogens depends on the infection window showing host mortality due to infection ($\zeta$), host generation time ($G_t$), and a scaling coefficient $f$. We modelled the lag period $g^{lag}$ based on the incubation period ($\psi$), where the pathogen proliferates to cause the acute infection phase and the host mortality window ($\zeta$) scaled by coefficients $l$ and $m$, respectively. The coefficients $l$ and $m$ also depend on generation time $G_t$, as described in Table S3. Note that, for the sake of simplicity, we considered the same values for $S_0$ and $S_{max}$ for all four pathogens with varying growth and clearance dynamics.

(b) Adaptation against coinfecting pathogens:

We formulated $g^{lag}$ and $\theta$ as:

$$\theta = \zeta_R.\zeta_{co}^{-1}$$

$$g^{lag} = (g_R^{lag} + \eta.\zeta_R.\zeta_S^{-1})$$

As described in the main text, we described parameter $\theta$ as the ratio of the mortality window after infection with the fast-growing vs coinfecting pathogens, denoted by $\zeta_R$ and $\zeta_{co}$, respectively, capturing the relative estimate of constraints imposed by fast-growing pathogens. Subsequently, we considered the lag period to adapt against coinfection ($g^{lag}$) as a summation of the predicted lag period to adapt against the rapidly growing pathogen ($g_R^{lag}$) and the ratio of the host mortality window against rapid- vs slow-growing pathogens (denoted by $\zeta_R$ and $\zeta_S$, respectively) along with a directionality coefficient ($\eta$). Overall, the parameters $\theta$ and $g_{co}^{lag}$ for variable coinfection scenarios influence the trajectory of host adaptation against coinfecting pathogens. We considered $\varepsilon = 0.2$ to obtain survival parameters for predicting host adaptation trajectories against coinfecting pathogens with low interference between them. We kept the $S_0$ and $S_{max}$ the same as the previous host adaptive trajectory prediction against individual pathogens. We simulated all the parameters in R.

**B. Baseline population and generation of experimental beetle populations**

We used a large outbred population (~2500 individuals) of *Tribolium castaneum* as the baseline stock population, maintained on whole wheat flour at 33℃ under constant darkness, with a discrete generation cycle of ~40 days, for more than two years before commencing the experiments (See ref (*4*) for a detailed description of the beetle population). We allowed ~1000 baseline stock adults to oviposit in 400g wheat flour to generate beetles for all experiments, including initiating experimental evolution regimes (each replicate population was initiated separately). Following this, we removed the adults after 48 hours and allowed the offspring to develop for 3 weeks. We sexed the offspring at the pupal stage and isolated virgin beetles in wells of 96-well micro-plates (with ~25mg wheat flour/well) for 2 weeks to allow adult-eclosion and attaining sexual maturity. Since the pupal stage typically lasts 3–4 days, we obtained 10-day-old virgin adults for all experiments.

**C. Bacterial infection and its effects on beetle survival and reproduction in the baseline population**

We infected beetles following a previously published protocol (*5*), using entomopathogens *Bacillus thuringiensis* DSM 2046 (Bt) (*5*) and *Pseudomonas entomophila* L48 (Pe) (*6*). The primary cultures were initiated from glycerol stocks and grown overnight in Luria Broth at 30°C with constant shaking (150 rpm) in bacterial incubators. The next day, a secondary culture was initiated from the primary culture and allowed to reach 1 OD (measured at 600 nm). We next centrifuged and resuspended the bacterial pellet in sterile insect Ringer solution to prepare the bacterial slurry and adjusted the final infection dose for Bt and Pe to 30OD (~ $8\times10^{\wedge7}$ cells/µl) and 100OD (~ $4\times10^{\wedge9}$ cells/µl). To prepare a mixed infection dose (Mx), we combined an equal proportion (1:1) of the resuspended culture of Bt and Pe cells. We used the same infection dose for all the experiments, including the infection treatments for experimental evolution regimes. To estimate the number of bacterial cells delivered inside the beetles, we infected 15-day-old larvae using the septic injury method (*5*), followed by plating the whole-body homogenate immediately (i.e., zero-hour bacterial load) on a Luria agar plate (n=10 larvae/infection treatment). While infecting with Bt and Pe yielded ~120 ± 7 cells/individual and ~ $2\times 10^4$ ± 500 cells/individual, respectively, mixed infection delivered ~30 ± 2 Bt cells + $1\times 10^3$ ± 500 Pe cells/ individual. Note that although we assayed adult beetles in all our experiments, we could not use their whole-body homogenate while estimating the zero-hour bacterial load. Whole-body extraction would have released the antimicrobial quinones from stink glands, which can kill the bacterial cells and thereby interfere with accurately estimating the total number of bacterial cells delivered inside the beetles during each infection (*7*, *8*). We can measure the bacterial load from adult beetles only after establishing a systemic infection (i.e., the spread of infection from the injection site, after ~3h post-infection) (described later). We pricked the beetles with sterile insect Ringer solution as a procedural control (sham infection) (*5*).

To estimate post-infection survival, we pricked 10-day-old (post-eclosion) adult virgin females derived from baseline populations between their head and thorax, using a 0.1 mm minute pin (Fine science tools) dipped in the bacterial suspensions (or sterile insect Ringer solution for sham infection). Next, we distributed them individually into 96-well plates with approximately 0.5 grams of fresh flour and monitored their survival daily for the next ten days (n=30 females/infection treatment). We analysed the survival data as a function of infection treatment using a Cox proportional hazard analysis (*9*).

Effects of infection on reproductive output

We also quantified the reproductive output of the infected beetles from the baseline population. To this end, we first obtained 10-day-old virgin males and females from our baseline population and infected (or sham-infected) them following the protocol described above. After 6hpi (so that beetles recover from the infection handling), we paired the males and females from each treatment with 1 g of doubly-sifted wheat flour (to remove bigger wheat flakes) in 35 mm Petri plates. We allowed mating and oviposition simultaneously for the next 36 hours (i.e., until 42hpi). Subsequently, the mating pairs were transferred to another plate supplemented with fresh flour, and the eggs were counted by sieving the sifted wheat through a 350-micron mesh (*10*). After another 36 hours (i.e., 78hpi), we subjected the surviving mating pairs to the second round of egg-laying for 36 hours and counted the number of eggs at 114hpi. We chose the first and second oviposition windows to capture the effects on reproduction during (i) the early infection phase, where mortality begins in all the infection treatments, and (ii) the later infection phase, when Bt cells are removed from the body of surviving individuals, but Pe cells persist (n= 30 mating pairs/infection treatment/oviposition window). The later infection phase also overlapped with the beginning of the reproductive window for our selection regimes (see the main text). We analysed the data using a generalized linear model fitted to the Poisson distribution with treatment as fixed effects. Pairwise contrasts between treatments were measured using "emmeans" adjusted with Tukey's HSD.

**D. Quantifying bacterial load dynamics after infection with single vs coinfecting pathogens in baseline populations**

Bacterial dynamics in live beetles after Bt, Pe and Mx infection

To measure the within-host bacterial growth dynamics, we subjected 10-day-old virgin females to respective infection treatments (i.e., Bt, Pe or Mx), followed by sampling of a subset of surviving beetles at regular intervals post-infection (Bt: every 2–4h until 24 hours; Pe: every 6–24h until 82 hours; Mx: every 2–24h until 82 hours). To this end, we followed a previously established protocol (10), where at each time point, we used micro-scissors (Fine Science tools, CA, USA) to first carefully remove quinone-containing thoracic and abdominal stink glands from each beetle, followed by pooling and homogenising the remaining carcass of 3 beetles in 60µl of Luria broth (LB) (stock homogenate). We then added this stock homogenate into a 96-well plate and made serial dilutions of the homogenate till $10^{-5}$-fold (i.e., $10^{-1}$–$10^{-5}$). We took 4µl of the homogenate of each serial dilution, spotted it on a Luria Agar plate, and incubated it overnight at 30°C (n=8–10 replicates of pooled homogenate of 3 females/infection treatment/timepoint). We also spread plated 20µl of the homogenate till $10^{-2}$ fold serial dilution (including the stock homogenate) using glass beads for Mx infection. The next day, we noted the colony-forming units (CFUs) from the minimum dilution of the homogenate that produced isolated colonies, followed by accounting for the appropriate dilution factor to estimate the bacterial load per beetle as described in Prakash et al. (*7*). We distinguished Bt and Pe cells for beetles infected with Mx by their colony morphology and size (see **Fig. S13** for representative colony morphology, validated separately by 16s rRNA sequencing).

In separate experiments, we also tracked the Pe cells in live beetles (a) after Pe infection at several time points between 4 and 25 days post-infection to establish the persistence of Pe cells in the host body and (b) on day 10 after Mx infection to show the presence of Pe infection even after the reproductive window is passed.

We analysed these log-transformed bacterial load data fitted to a gamma distribution (based on the lowest Akaike information criterion (AIC) value), with treatment and time of assay as fixed effects. To allow the inclusion of replicates with zero bacterial load in the analysis, we added two CFUs to all bacterial load data before log transformation.

Bacterial load in dead beetles after Mx infection

Separately, we measured bacterial load from individuals who died after infection with Mx at various time points until 46 hours (n=34 females). We individually crushed the carcass obtained from dead females (as described above) in 60µl of chilled LB. Since bacterial load in dead individuals could be higher than in alive individuals, we made serial dilutions till $10^{-7}$ and plated all the dilutions. We ensured that dead beetles were dissected and plated immediately to avoid excessive bacterial growth in the cadaver without any active immune system.

**E. Quantifying within-host bacterial load and growth dynamics in evolved beetles**

Bacterial load estimation at the onset of mortality:

To test whether the increased survival of standardised B- and P-beetles (tested at generation 22 and 18, respectively; see the main text) can be explained by their improved ability to clear pathogens, we measured the bacterial load across replicate populations at the onset of the first 10–15% mortality after the respective infection treatments: e.g., Bt: 8hpi and Pe: 24hpi (n=10–15 replicates with pooled homogenate of 3 females/selection regime/replicate population). We checked the bacterial load for standardised M-beetles (tested at generation 18) at 8 and 20 hours to capture adequate Bt and Pe cells, respectively (n=10–15 replicates with pooled homogenate of 3 females/replicate population/time point). For each pathogen-selected regime, we compared the bacterial load (Bt and Pe cells analyzed separately in case of Mx infection) at these time points with respect to the load obtained from females from the control regime after the respective infection treatments. We analysed the bacterial load for each selection regime with respect to C-beetles, using the generalized linear mixed effects model (using the "glmer" function) fitted to Gamma distribution, with the selection regime as a fixed effect and replicate population as a random effect.

Temporal dynamics of bacterial load across section regimes

We followed the above protocol to measure the temporal changes in bacterial load in the standardised beetles from the selected vs. control regime in generation 25 (for Mx infection) and generation 26 (for Bt and Pe infection). We sampled infected alive females at multiple time points based on their with-host growth dynamics patterns observed in our previous experiments. For instance, since Bt could grow fast, followed by rapid clearance by the beetle host, we estimated the bacterial load of B-beetles every few hours until 18hpi, whereas P-beetles infected with Pe causing long-lasting infection were assayed until 72hpi (n=6–10 replicates with pooled homogenate of 3 females/selected regime/time point). We measured the bacterial load of M beetles every few hours until 50 hours to detect both Bt and Pe cells (n=6–10 replicates with pooled homogenate of 3 females/selected regime/time point). We also noted the bacterial load from every individual who died after infection with Bt, Pe and Mx across selection regimes during the first 12 hours, 30 hours and 50 hours respectively. We separately analysed the bacterial load data of (a) alive beetles for each pathogen and regime, using a generalized linear model fitted to Gamma distribution, with selection regime and time points as fixed effects; (b) dead beetles using a generalized linear model fitted to Gamma distribution, with the selection regime as a fixed effect. Note that due to logistical challenges, we could only assay replicate population 3 from each selection regime.

We also studied the growth and clearance of Bt cells more closely during the early coinfection phase by measuring the bacterial load every hour between 6 and 12hpi in live M- vs C-beetles from replicate population 4 in generation 30. This allowed us to broadly compare the bacterial growth dynamics patterns during the early infection phase observed in replicate population 3. We analysed the data as described above.

**F. Validation of beetle mortalities caused by Bt growth during the early coinfection phase across all the replicate populations M- vs C-beetles**

We also quantified the number of beetles that died in all M- vs C-regime replicate populations within the first 20 hours of Mx infection (n= 30 females/selected regime/replicate population; tested at generation 28) and validated if they carried an excess of Bt load upon death as described earlier (~$10^5$ Bt cells/ beetles). We analysed the (a) total number of beetle mortality across replicate populations and selection regimes, using a pairwise Wilcoxon test; (b) bacterial load data using a generalized linear model fitted to Gamma distribution, with selection regime as a fixed effect.

**G. Transcriptomic analysis:**

We passed the raw reads through *FastQC* and *MultiQC* to check the quality parameter summaries. We then used *Cutadapt* to remove adapter contamination using standard Illumina adapter sequences (forward adapter sequence: AGATCGGAAGAGCACACGTCTGAACTCCAGTCA and reverse adapter sequence: AGATCGGAAGAGCGTCGTGTAGGGAAAGAGTGT along with AAAAAAAAAA on both sides for poly-A tail removal). During adapter trimming, we discarded reads with N base counts more than 10% of its length and more than 10 low-quality bases, and restricted the final read lengths ranging between 70–150 base pairs for the subsequent analyses. We then made a genomic index using the *T. castaneum* reference genome Tcas5.2 (GCF_000002335.3) (*11*) and mapped all the reads using *Hisat2,* allowing for up to 3 mismatches per read along with strand information. Subsequently, we used *stringtie 2* for sorting bam output files.

We further quantified reads using *HTseq 2.0.3* by restricting read alignment by positions and implemented a strict option for intersectional reads along with reverse strand-specific exon mapping. Statistical analyses were done in R 4.1.3, with the package "DESeq2". Subsequently, we filtered for genes having more than 1 normalized read count in each sample and excluded the rest for further analyses, resulting in a total of 18,076 genes. The raw read counts were then normalized, and normalized log2 count values were further analysed to estimate differential gene expression. We used linear models within *DESeq2,* taking each combination of selection regime and infection treatment, and replicate population as fixed effects (normalized $\log_2$ count ~ infection treatments+ replicate population) to calculate log-fold changes in gene expression. Since replicate populations were handled independently and might have different adaptive trajectories, we estimated paired contrasts between each replicate population across combinations of selection regime and infection treatment. Here, we have considered differential expression between each infected population and their sham-infected counterparts across the control and three pathogen-selected regimes (n=4 replicates; each comprised of 10 females pooled together from each replicate population/infection treatment/selection regime). We have filtered differentially expressed genes (DEGs) based on P-value <0.001. We visualized expression profiles of all the DEGs by a pooled-population heatmap based on the z-score of normalized read counts using "pheatmap" and "RColorBrewer" in R.

We further performed a principal component analysis based on the normalized count data of all the DEGs from all three pathogen-selected regimes. We have also estimated the common and unique set of differentially regulated DEGs, including up- and down-regulated genes, among the three pathogen-selected regimes and visualized them using the R-package “ComplexUpset”. Subsequently, we used *Blast2Go* to find all the gene ontology terms (GO terms). We further performed pathway enrichment with the GO terms for KEGG, using the R-package “gProfiler2” and plotted enriched pathways using R-package “ggplot2”. We also manually compiled all the known immunity-related DEGs and categorized them under five main classes based on the previous literature: (1) pathogen receptors and immune pathway receptors (e.g. Toll and Imd pathway, Jak-Stat pathways), (2) regulator molecules and intracellular signal transducers, (3) inducible immune effector molecules such as antimicrobial peptides and lysozyme, (4) phenoloxidase mediated melanization response and (5) reactive oxygen species-mediated response.

Subsequently, we estimated a gene expression profile using generalized canonical discriminant analyses implementing linear models (LDA) with each category of immune-related DEGs separately by implementing the R-package “canDisc” as described in Berger et al. (*12*). We always used normalized count values of these categorized DEGs. We only considered the first linear discriminant (LD) axis as the gene expression profile estimate to avoid overfitting. We performed Tukey’s HSD to obtain the differences in immune responses between control and selected populations after the respective infection treatments. To understand the association between expression changes of each immune gene category with variations in phenotype we performed multiple regression and canonical correlation analysis. At first, we performed linear multiple regression analyses between gene expression profile and the hazard ratios (as a proxy of survival response estimated from the survival data of each replicate population across selection regimes after the respective infection treatment vs sham infection) or bacterial load individually. Next, we implemented canonical correlation analyses (CCA) using the R-package “canDisc” between gene expression profile and combined phenotypic profile (comprising both hazard ratio and bacterial load for each infection treatment and selection regime) to examine how differences in gene expression correlate with the phenotypic variations. In CCA, we took the first three LD axes as three x-variables representing cumulative gene expression profiles for each immune gene category. Subsequently, we used hazard ratios and their bacterial load (log CFU) values as two y-variables, representing the phenotypic variations. We then estimated the canonical R-squared values between each immune gene category and phenotypic variations for each combination of pathogen-selected and their respective control populations upon infection (i.e. B and C regime infected with Bt; M and C regime infected with Mx; P and C regime infected with Pe). To visualize the correlation space, we took the first canonical axis for both selected and control populations and performed linear regression across all three pathogen-selected regimes.

#### H. Validating the gene expression profile of phenoloxidase response in B- vs C- regimes

We followed our previously published protocol to measure the enzymatic activity of phenoloxidase (PO) response as described in Khan et al. (*13*). Since PO response can be induced rapidly (i.e., as early as 1hpi (*14*) we infected standardized females from B and C regimes with Bt (or sham infection with Ringer solution) and assayed the PO activity of individual beetles within 3 hours post-infection (n= 9–12 females/infection treatment/regime/replicate population). Note that we experimented with only two replicate populations. We separately analysed each replicate population across selection regimes, using the Wilcoxon rank sum test as a function of infection status.

#### I. Packages and software used for visualization:

We used R statistical programming language (version 4.1.3, R Development Core Team, 2019) for carrying out the different statistical tests and visualization of the data. The following packages were used for visualization: “ggplot2”, “ggkm”, “tidyr”, “dplyr”, “ggpubr”, “patchwork”, “survival” and “survminer”. Adobe Illustrator (2023) was used for compiling the multi-panel figures mentioned in the manuscript. Figure 1 was designed using the Biorender platform.

**Supplementary figures**

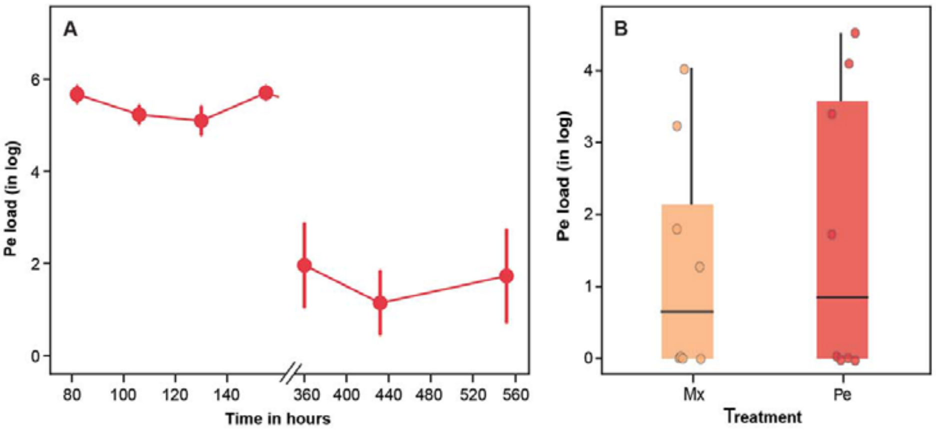


**Fig. S1:** (**A**) Persistence of *P. entomophila* (Pe) cells until 22 days post-infection (n= 4–5 replicates with pooled homogenate of 3 females/time point). (**B**) The number of Pe cells in beetles on the 10th day after infection with Mx vs Pe (n= 8 replicates with pooled homogenate of 3 females/time point).

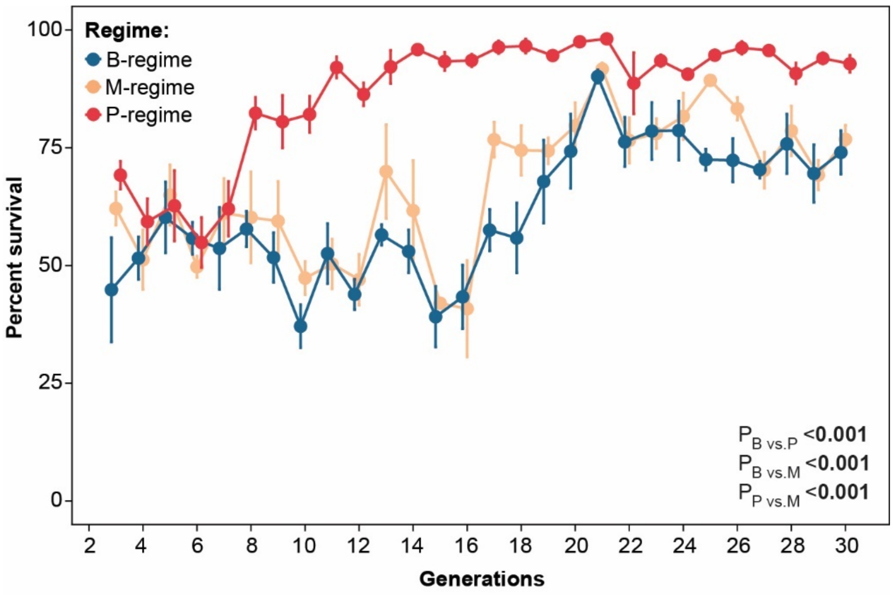


**Fig. S2:** Beetle survival across generations (Generation 3–30) at the beginning of the oviposition window (i.e., 3rd day post-infection) in each replicate population of different pathogen-selection regimes (n= 4 replicate populations/selection regime). P-values represent the pairwise differences between selection regimes.

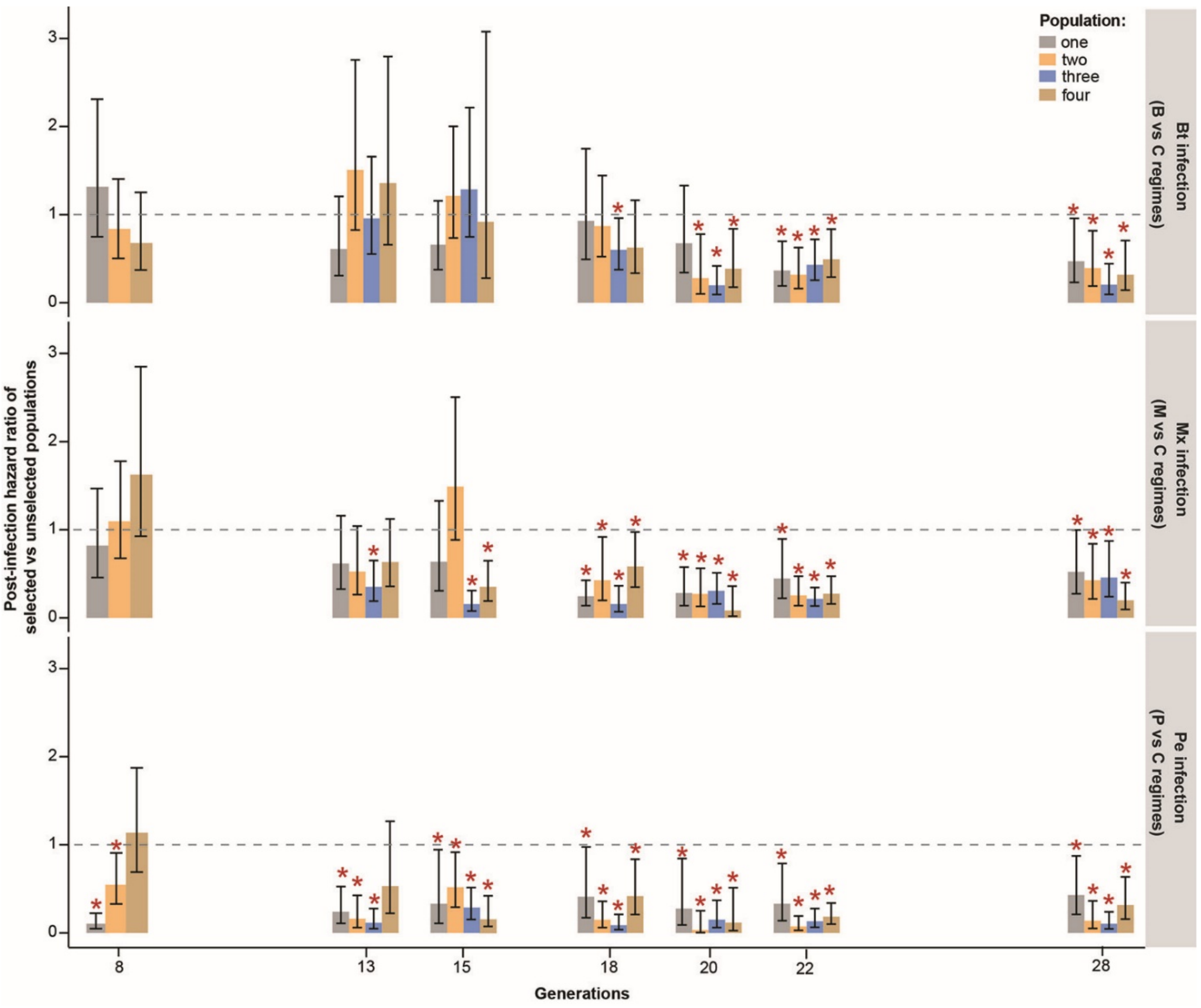


**Fig. S3: An estimate of survival response in each replicate population of pathogen selected (i.e., P-/ B-/M-regime) vs control beetles after the respective infection treatments (e.g., Pe/Bt/Mx) at different generations during the experimental evolution.** The dashed lines indicate a hazard ratio of 1 (i.e., no change). A hazard ratio of less than 1 indicates the evolution of increased post-infection survival. Asterisks denote hazard ratios significantly less than 1 (Cox proportional hazard analyses, $p<0.05$). Every generation, we assayed all 4 replicate populations (except at generation 8, where only 3 populations were assayed) (n= 24–60 females/replicate population/selection regime).

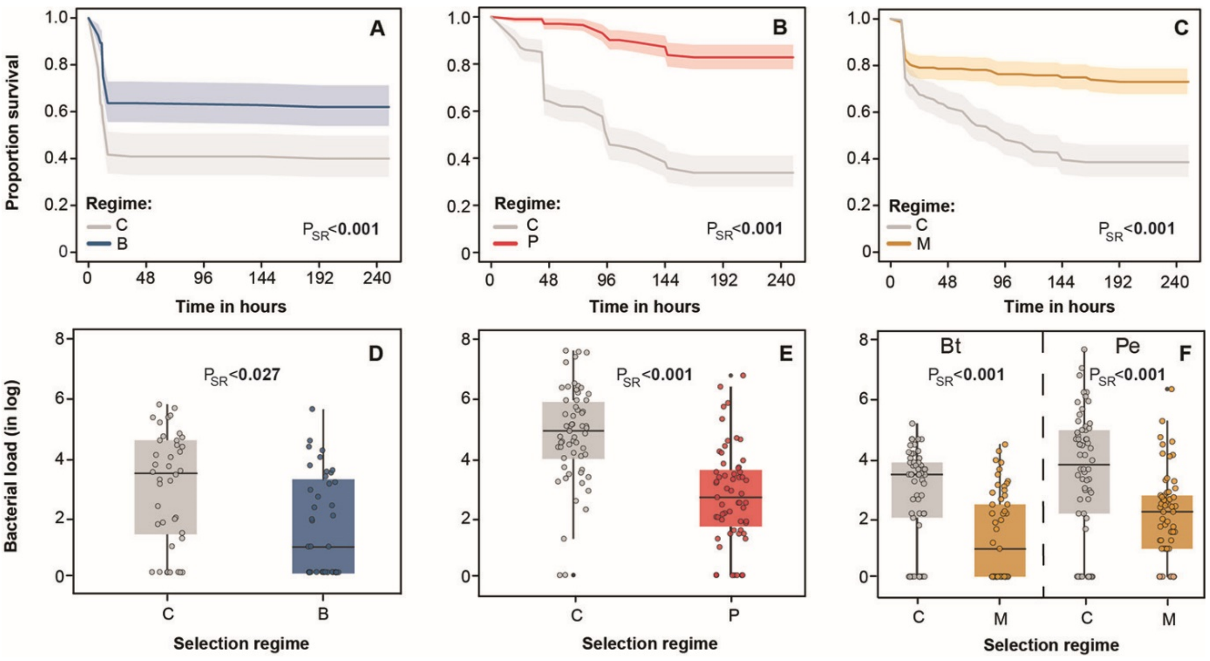


**Fig. S4: Correlating increased post-infection survival in selected regimes with lower bacterial load.** Survival of virgin females from the selected vs control regimes following infection with (**A**) Bt, (**B**) Pe, and (**C**) Mx (n=32 females/replicate population/selection regime) (Data analysed using mixed effect Cox model). Bacterial load after (**D**) Bt; (**E**) Pe and (**F**) Mx infection in the selection vs control females (n= 10–15 replicates with pooled homogenate of 3 females/replicate population/selection regime) (Data analysed using generalized linear mixed model fitted to Gamma distribution). P-values indicate the main effects of the selection regime.

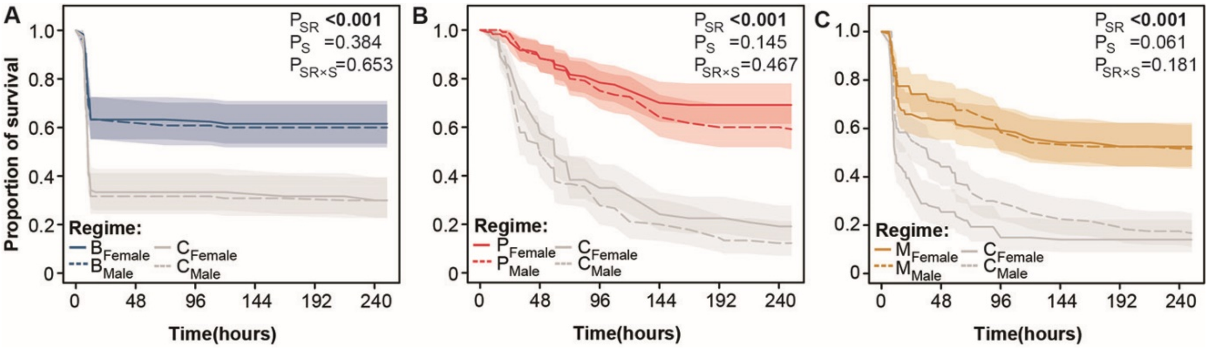


**Fig. S5: Comparing the increase in post-infection survival across sexes.** Post-infection survival in virgin males and females from selected (i.e., P/ B/ M) - vs control (C) regimes following infection with **(A)** Bt; **(B)** Pe; and **(C)** Mix (n= 30 beetles/sex/replicate population/selection regime) (Data analysed using mixed effect Cox model). P-values indicate the main effects of the selection regimes, sex and their interaction with significant values highlighted in bold.

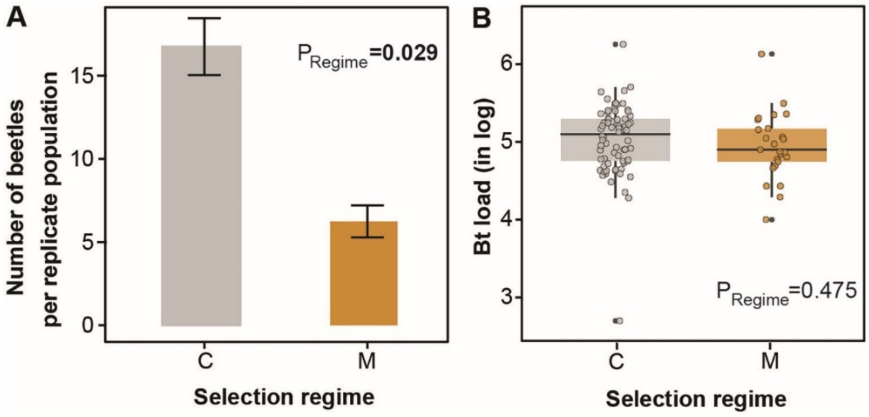


**Fig. S6: (A)** The total number of individuals that succumbed to Mx infection in C- vs M-regime within the first ~20 hours post-infection (n= 5–7 females/replicate population from M-regime; n=15–19 females/replicate population from C-regime) (data analysed using Wilcoxon rank sum test) and (**B**) their *B. thuringiensis* (Bt) load (data analysed using generalized linear mixed model fitted to Gamma distribution). P-values indicate the main effects of the selection regimes with significant values highlighted in bold.

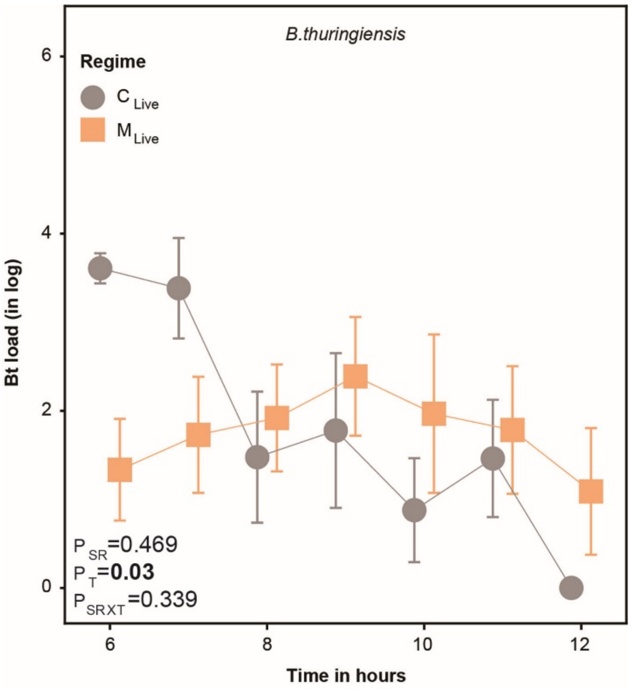


**Fig. S7: Temporal changes in the clearance of Bt cells across M- and C-regime after Mx infection**. Bt load was measured every hour between 6- and 12hpi (n= 6–8 replicates with pooled homogenate of 3 females/time point/selection regime) (data between 8–12 hours was analysed using generalized linear model fitted to Gamma distribution). P-values indicate the main effects of the selection regimes, time points and their interactions with significant values highlighted in bold.

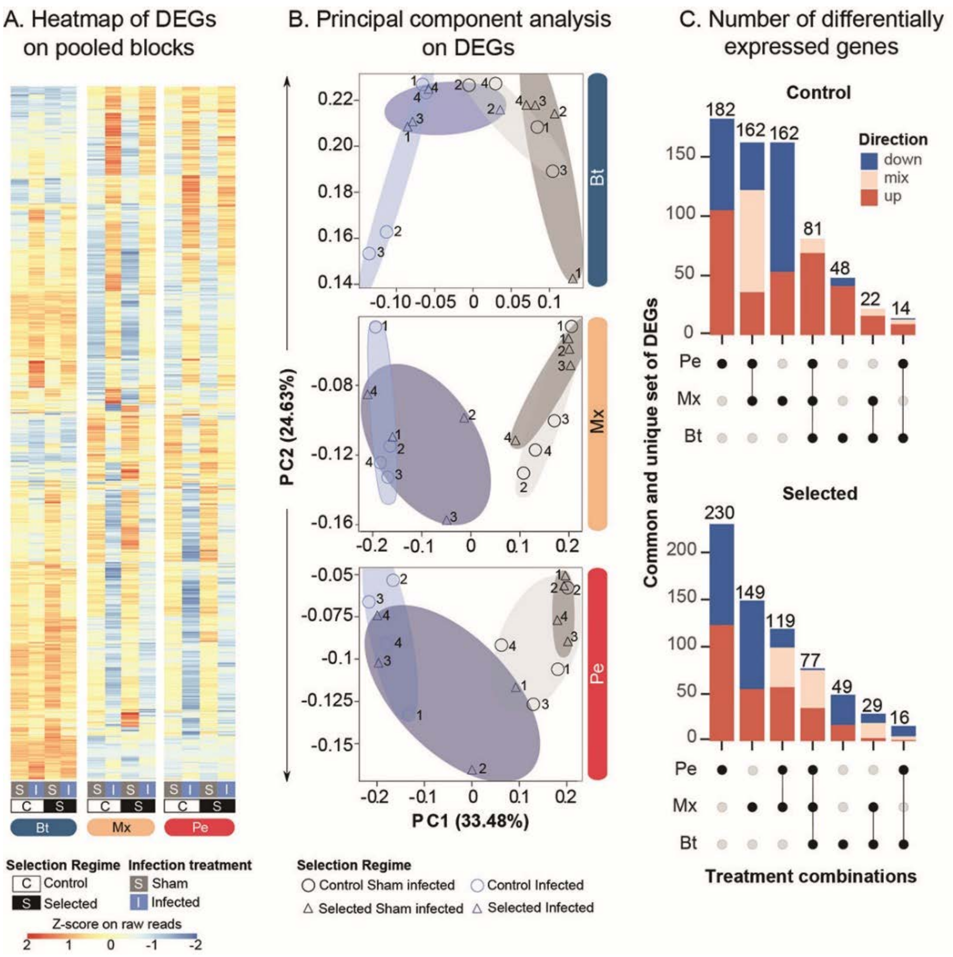


**Fig. S8: Differentially regulated genes after experimental evolution.** (**A**) Heatmap based on Z-scores of normalized read counts of differentially expressed genes (n ~700) with infection (i.e., Bt, Pe and Mx infection) in control vs selected regimes. (**B**) Principal component analysis on differentially expressed genes in each of the pathogen-selected regimes (B, P or M) and corresponding C beetles after their respective infection treatments with Bt, Pe and Mx. Blue shades denote infection treatments, whereas grey shades denote sham treatments. Numbers correspond to each replicate population. (**C**) Upset plot showing the total number of differentially expressed genes (DEGs): upregulated, downregulated, or with contrasting expression patterns due to infection in control vs pathogen-selected regimes after their respective infection treatments.

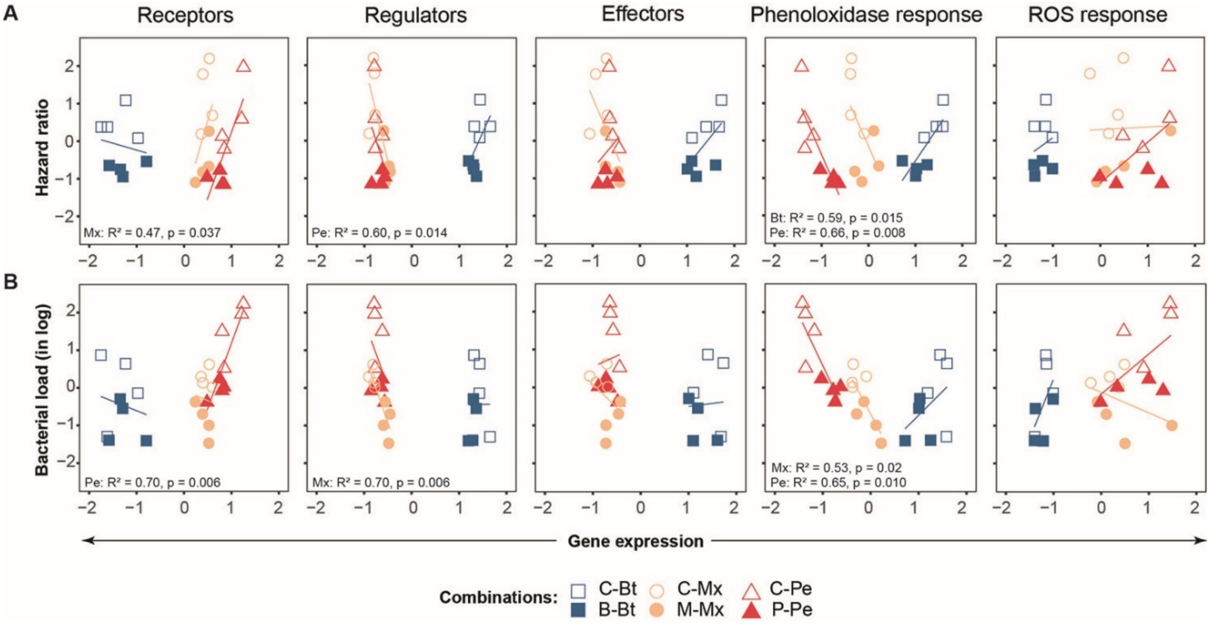


**Fig. S9: Correlation between changes in gene expression profiles of immune genes and phenotypic traits such as hazard ratio and bacterial load.** A linear regression analysis to show the association of gene expression profile with hazard ratio (**A**) and bacterial load (**B**) in unselected control and corresponding selected regime post-infection with either Bt, Pe or Mx. The first axis of LDA (**Fig 5B**) is considered as gene expression profile for various categories of differentially expressed immune-related genes, namely (a) pathogen and immune receptors; (b) immune regulators; (c) inducible immune effectors, including antimicrobial peptides (AMPs) and lysozymes; (d) fast-acting phenoloxidase mediated melanisation response; and (e) production of reactive oxygen species. In each panel, regression statistics for each significantly associated group are provided (see **Table S24,25**).

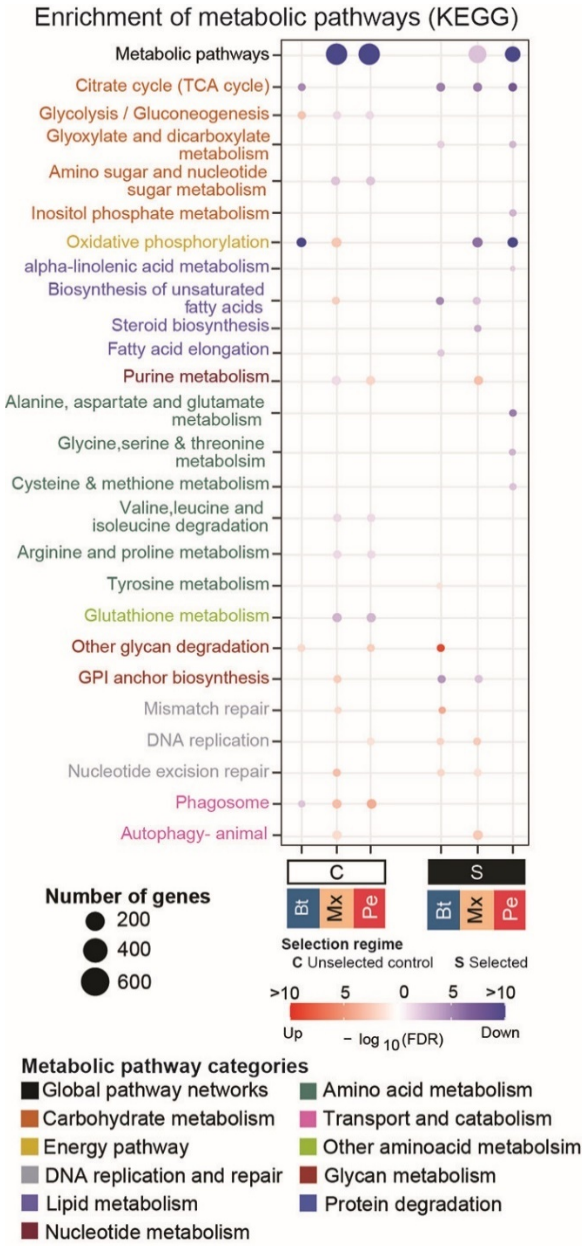


**Fig. S10: Effects of infection and pathogen selection on metabolism.** Enrichment plot indicating the number of metabolic pathway components that are either differentially upregulated or downregulated in control vs selected populations post-infection. The pathways are colour-coded based on the classifications provided in KEGG. In each of the contrasts in the control vs selected regime across pathogens, changes in expression level post-infection were obtained by comparing them with respective sham infections.

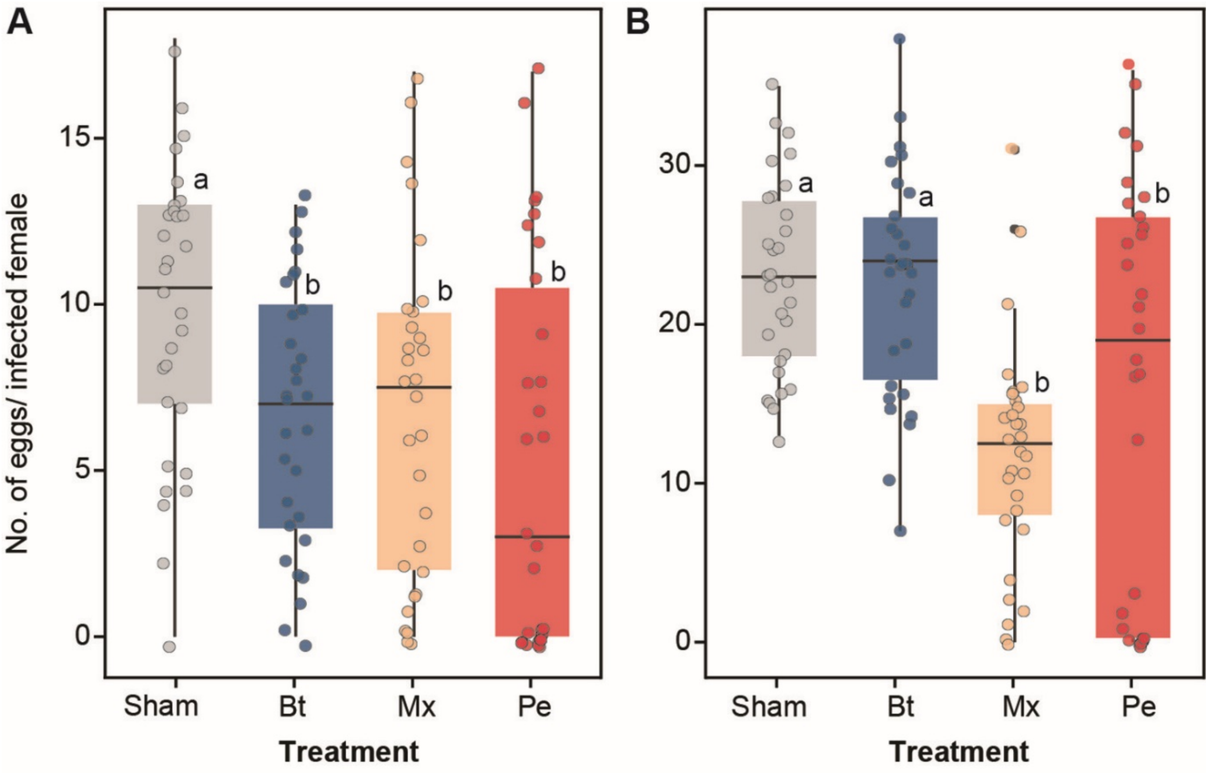


**Fig. S11: Effect of bacterial infection on fecundity at different time points post-infection.** The number of eggs laid by females after Bt, Pe, Mx or Sham infection (pricked with sterile Ringer solution) (A) between 6 and 42hpi (B) between 78 and 114hpi (which corresponds to the beginning of oviposition window for the selection regimes; also see methods). (n= 30 mating pairs/infection treatment/oviposition window) (data analysed using generalized linear model fitted to Poisson distribution). In each panel, significantly different groups are connected with different alphabets (based on Tukey's HSD).

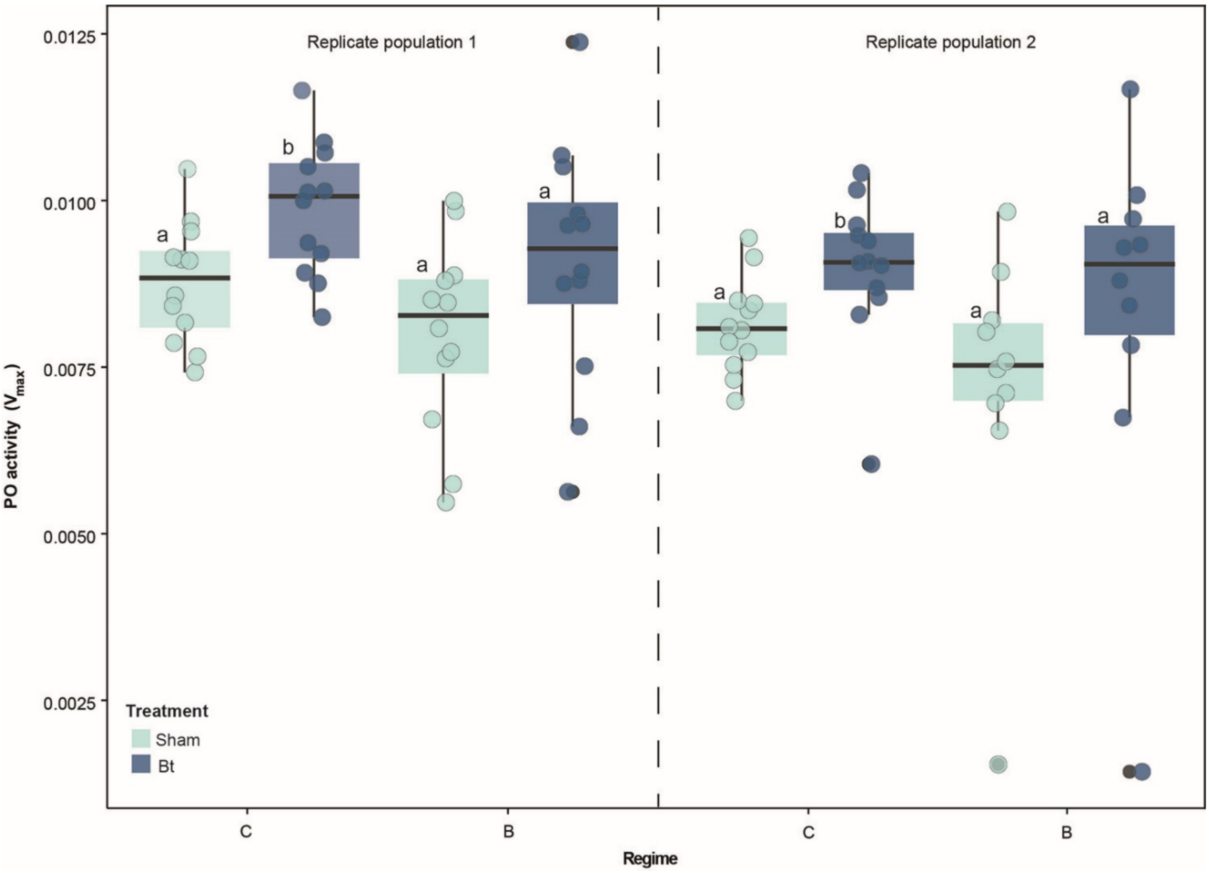


**Fig. S12: Comparing the phenoloxidase activity after Bt infection in B- vs C-regime.** PO activity was measured in replicate populations 1 and 2 from B- and C-regimes as Vmax of the enzymatic reaction (slope of the linear phase of the reaction curve) (n= 8–10 females/infection treatment/selection regime/replicate population) (data analysed using Wilcoxon rank sum test). In each regime, significantly different groups (infection treatments) are connected with different alphabets and are not comparable between regimes.

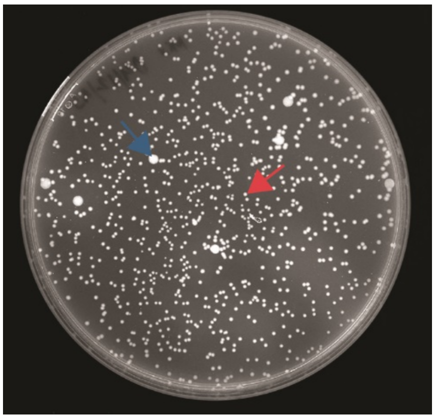

**Fig. S13: Representative colony-forming units of Bt and Pe on a Luria Agar plate.** The bacterial cells were obtained from plating pooled homogenate of three females post-infection with Mx. A single colony of Bt and Pe is denoted by blue and red arrows respectively.

### SUPPLEMENTARY TABLES

**Table S1:** The table provides a summary of combined demographic model parameters, as described in Dunaeu et al. 2017 (*1*). These parameter values were used to simulate four divergent within-host growth dynamics, such as rapidly growing pathogens causing acute infections, followed by (1) rapid clearance ($R_C$) or (2) persistent infection ($R_P$); Slow-growing pathogens causing acute infection, followed by (3) rapid clearance ($S_C$) or (4) persistent infection ($S_P$).

| Parameters | Symbols | Pathogens with different growth dynamics | | | |
|---|---|---|---|---|---|
| | | **Rc** | **Sc** | **Rp** | **Sp** |
| *Baranyi Model* | | | | | |
| 1. Bacterial load upon injection | $n_0$ | 1 | 3 | 2 | 3 |
| 2. Maximum bacterial load | $n_{max}$ | 6 | 10 | 6 | 10 |
| 3. Lag time to growth | $t^{lag}$ | 4 | 8 | 1 | 4 |
| 4. Early bacterial growth rate | $\mu$ | 0.9 | 0.4 | 0.7 | 0.3 |
| 5. Standard deviation | $\sigma^b$ | 0.035 | 0.037 | 0.31 | 0.36 |
| *Control and survival* | | | | | |
| 6. Time of control | $t_C$ | 6–7 | 13–14 | 3–4 | 7–8 |
| 7. Proportion of survival | $P_S$ | 0.41 | 0.38 | 0.32 | 0.29 |
| *Exponential decline model* | | | | | |
| 8. Decrease rate in bacterial load when infection is controlled | $\delta$ | 1 | 0.4 | 0.02 | 0.02 |
| 9. Standard deviation | $\sigma^d$ | 0.033 | 0.035 | 0.036 | 0.038 |

**Table S2:** The table provides a summary of the parameters used to simulate survival patterns against individual pathogens with divergent growth dynamics: (a) post-infection incubation window (i.e., a period after infection with no host mortality) and (b) the mortality window (i.e., a period that encompasses all the host mortality due to infection).

| Parameters | Symbols | Pathogens with divergent growth dynamics | | | |
|---|---|---|---|---|---|
| | | **Rc** | **Sc** | **Rp** | **Sp** |
| 1. Post-infection incubation window | $\psi$ | 7 | 14 | 4 | 7 |
| 2. Mortality window | $\zeta$ | 5 | 6 | 16 | 14 |

**Table S3:** The table provides a summary of parameters for predicting host adaptation against individual pathogens vs. coinfections.

| Parameters | Symbols | Values for adaptation trajectory simulation |
|---|---|---|
| 1. Interference coefficient | $\varepsilon_{RS}$, $\varepsilon_{SR}$ | 0.2 |
| 2. Host survival upon infection in ancestral population | $S_0$ | 0.2 |
| 3. Host survival upon infection in evolved population | $S_{max}$ | 0.9 |
| 4. Generation time | $G_t$ | 30 × 24 |
| 5. Scaling factor for mortality window | $m$ | $(G_t/2)^2$ |
| 6. Scaling factor for incubation period | $I$ | $(G_t/6)^2$ |
| 7. Directionality coefficient | $\eta$ | $(\zeta_R - \zeta_S)/(\lvert\zeta_R - \zeta_S\rvert)$ |

**Table S4: Response to single vs coinfection in baseline stock population.** The table provides a summary of Cox proportional hazard analyses of survival data after infection with single pathogen (Bt or Pe) or coinfection (Mx) [Model specification: Survival ~ Infection treatment]. Bt, Pe and Mx denote infection with *B. thuringiensis*, *P. entomophila* and a mix of both Bt and Pe, respectively.

| Effects | Loglikelihood | df | $\chi^2$ | P |
|---|---|---|---|---|
| Across different infection types (i.e., Bt, Pe & Mx) | -229.50 | 2 | 0.1471 | 0.92 |
| Pairwise comparison | | | | |
| Bt vs Mx | 127.37 | 1 | 0.2121 | 0.64 |
| Bt vs Pe | 140.09 | 1 | 0.1377 | 0.71 |
| Pe vs Mx | 146.22 | 1 | 0.1483 | 0.70 |

**Table S5: Analysis of Bt and Pe load dynamics during single-infection vs coinfection scenario in baseline stock population**. The table provides a summary of a generalised linear model on log-transformed bacterial load data of (**a**) Bt and (**b**) Pe cells separately, fitted to a Gamma distribution [Model specification: Bacterial load ~ Infection type (I) (i.e., Single vs coinfection) + Time points (T) + Infection type × Time points]. Significant effects are highlighted in bold.

| Pathogen | Effects | df | $\chi^2$ | P |
|---|---|---|---|---|
| **(a)** Bt | I | 1 | 19.44 | **<0.001** |
| | T | 5 | 501.48 | **<0.001** |
| | I × T | 5 | 32.32 | **<0.001** |
| **(b)** Pe | I | 1 | 0.003 | 0.9560 |
| | T | 6 | 38.459 | **<0.001** |
| | I × T | 6 | 12.570 | **0.0503** |

**Table S6: Changes in percentage survival in the pathogen-selected regimes (i.e., B, P, and M) across generations**. The table shows the summary of a generalised linear model on the percentage of beetles surviving every generation after three and eight days post-infection fitted to Gaussian distribution [Model specification: Survival ~ Selection regime (SR) + Generations (G) + Selection regime × Generations]. Pairwise contrasts between B-, P- and M-regimes were calculated using emmeans. Significant effects are highlighted in bold.

| Timepoint | Effects | $\chi^2$ | df | P |
|---|---|---|---|---|
| 3 days post-infection | SR | 388.27 | 2 | **<0.001** |
| | G | 422.27 | 27 | **<0.001** |
| | SR × G | 163.50 | 54 | **<0.001** |
| 8 days post-infection | SR | 185.39 | 2 | **<0.001** |
| | G | 586.12 | 27 | **<0.001** |
| | SR × G | 171.12 | 54 | **<0.001** |

Pairwise comparisons

| Timepoint | Contrasts | df | T.ratio | P |
|---|---|---|---|---|
| 3 days post-infection | B vs. P | 252 | -18.726 | **<0.001** |
| | M vs. B | 252 | -4.051 | **<0.001** |
| | M vs. P | 252 | -14.675 | **<0.001** |
| 8 days post-infection | B vs. P | 252 | -12.261 | **<0.001** |
| | M vs. B | 252 | -1.002 | 0.576 |
| | M vs. P | 252 | -11.259 | **<0.001** |

**Table S7: Survival response across selection regimes. (a)** The table provides a summary of mixed effects Cox model on post-infection survival data of females from control vs. pathogen selected regimes (i.e., B vs C, P vs C and M vs C), with replicate populations as a random effect [Model specification: Post-infection survival ~ Selection regime (SR) +(1|Replicate population (RP)); **(b)** The table provides a summary of the Cox proportional hazard analysis on post-infection survival data of beetles from control vs. pathogen-selected regimes, with replicate populations as a fixed effect [Model specification: Post-infection survival ~ SR + RP + SR × RP]. Significant effects are highlighted in bold. **(c)** The table provides a summary of the estimated hazard ratio of selected beetles relative to the control beetles after their respective infection treatments, calculated from the mixed effects Cox model as described in (**a**). A hazard ratio significantly <1 denotes an improved post-infection survival of selected beetles than control beetles.

| Effects | Loglikelihood | $\chi^2$ | df | P |
|---|---|---|---|---|
| **(a)** With replicate populations as a random factor | | | | |
| SR (B vs. C) | -601.49 | 21.567 | 1 | **<0.001** |
| SR (P vs. C) | -904.27 | 127.27 | 1 | **<0.001** |
| SR (M vs. C) | -999.77 | 64.163 | 1 | **<0.001** |
| **(b)** With replicate populations as a fixed factor | | | | |
| 1. B vs. C | | | | |
| SR | -605.12 | 14.300 | 1 | **<0.001** |
| RP | -603.19 | 3.0585 | 3 | 0.119 |
| SR × RP | -603.51 | 0.1715 | 3 | 0.941 |
| 2. P vs. C | | | | |
| SR | -911.04 | 113.75 | 1 | **<0.001** |
| RP | -961.57 | 15.937 | 3 | **<0.001** |
| SR × RP | -902.99 | 0.1156 | 3 | 0.141 |
| 3. M vs. C | | | | |
| SR | -1010.1 | 43.439 | 1 | **<0.001** |
| RP | -1008.2 | 3.8740 | 3 | 0.05 |
| SR × RP | -1007.2 | 1.9875 | 3 | 0.15 |

**(c)** Estimated hazard ratio

| Regime | Hazard ratio | P | Lower 95% | Upper 95% |
|---|---|---|---|---|
| B vs. C | 0.4118 | **0.047** | 0.1647 | 1.030 |
| P vs. C | 0.1879 | **0.001** | 0.0679 | 0.520 |
| M vs. C | 0.2174 | **<0.001** | 0.0956 | 0.4942 |

**Table S8: Changes in bacterial load across selection regimes. (a)** The table provides a summary of generalized linear mixed model on log transformed bacterial load data of females from control vs. pathogen-selected regimes (i.e., B vs C, P vs C and M vs C) fitted to a Gamma distribution, with replicate populations as a random effect [Model specification: Bacterial load ~ Selection regime (SR) +(1|Replicate population (RP))]. **(b)** The table provides a summary of generalized linear model on log transformed bacterial load data of females from control vs. pathogen-selected regimes (i.e., B vs C, P vs C and M vs C) fitted to a Gamma distribution, with replicate populations as a fixed effect [Model specification: Bacterial load ~ SR + RP + SR × RP]. Significant effects are highlighted in bold.

| Effects | $\chi^2$ | df | P |
|---|---|---|---|
| **(a)** With replicate populations as a random factor | | | |
| (A) SR (B vs. C) | 8.9828 | 1 | **0.0027** |
| (B) SR (P vs. C) | 35.036 | 1 | **<0.001** |
| (C) SR (M vs. C) | | | |
| (i) Bt-load | 21.441 | 1 | **<0.001** |
| (ii) Pe-load | 18.755 | 1 | **<0.001** |
| **(b)** With replicate populations as a fixed factor | | | |
| (A) B vs. C | | | |
| SR | 5.3575 | 1 | **0.02** |
| RP | 1.9220 | 3 | 0.16 |
| SR × RP | 1.9908 | 3 | 0.15 |
| (B) P vs. C | | | |
| SR | 7.3944 | 1 | **0.006** |
| RP | 1.9173 | 3 | 0.17 |
| SR × RP | 0.5674 | 3 | 0.57 |
| (C) M vs. C | | | |
| (i) Bt-load | | | |
| SR | 4.5825 | 1 | **0.03** |
| RP | 0.4384 | 3 | 0.51 |
| SR × RP | 0.2260 | 3 | 0.6345 |
| (ii) Pe-load | | | |
| SR | 10.619 | 1 | **0.001** |
| RP | 0.0794 | 3 | 0.78 |
| SR × RP | 3.2537 | 3 | 0.07 |

**Table S9: Response to selection across sexes**. The table provides a summary of mixed effect Cox model on post-infection survival data, with selection regime (SR) and sex (S) as fixed effects and replicate populations (RP) as a random effect [Model specification: Post-infection survival ~ SR +S+SR × S + (1|RP)]. Significant effects are highlighted in bold.

| Infection | Effects | Loglikelihood | $\chi^2$ | df | P |
|---|---|---|---|---|---|
| Bt | SR | -1510.7 | 58.586 | 1 | **<0.001** |
| | S | -1541.6 | 0.756 | 1 | 0.384 |
| | SR × S | -1510.1 | 0.203 | 1 | 0.653 |
| Pe | SR | -1573.9 | 113.267 | 1 | **<0.001** |
| | S | -1636.7 | 2.123 | 1 | 0.145 |
| | SR × S | -1572.7 | 0.525 | 1 | 0.467 |
| Mx | SR | -1727.9 | 81.237 | 1 | **<0.001** |
| | S | -1769 | 3.511 | 1 | 0.061 |
| | SR × S | -1725.3 | 1.788 | 1 | 0.181 |

**Table S10: Analyses of the temporal bacterial load dynamics in a representative population of selected vs. control beetles after the respective infection treatments.** The table provides a summary of a generalised linear model on log-transformed bacterial load data as a function of selection regime (SR) and time point (T), fitted to a Gamma distribution, from live [Model specification: Bacterial load ~ SR +T+SR × T] and dead beetles [Model specification: Bacterial load ~ SR] analysed separately. **(a)** C- and M-regime (i.e., C3 vs M3) after coinfection **(b)** C- and B-regime (i.e., C3 vs B3) after Bt-infection **(c)** C- and P-regime (i.e., C3 vs P3) after Pe-infection. Significant effects are highlighted in bold.

| Comparisons | Beetles | Bacteria | Effects | $\chi^2$ | df | P |
|---|---|---|---|---|---|---|
| **(a)** C vs M | *Live individuals* | Bt | SR | 2.643 | 1 | 0.104 |
| | | | T | 220.882 | 7 | **<0.001** |
| | | | SR × T | 13.812 | 7 | 0.054 |
| | | Pe | SR | 43.692 | 1 | **<0.001** |
| | | | T | 98.831 | 7 | **<0.001** |
| | | | SR × T | 33.254 | 7 | **<0.001** |
| | *Dead individuals* | Bt | SR | 0.6219 | 1 | 0.43 |
| | | Pe | SR | 28.123 | 1 | **<0.001** |
| **(b)** C vs B | *Live individuals* | Bt | SR | 1.241 | 1 | 0.2654 |
| | | | T | 159.026 | 4 | **<0.001** |
| | | | SR × T | 4.598 | 4 | 0.3310 |
| | *Dead individuals* | Bt | SR | 0.03 | 1 | 0.8618 |
| **(c)** C vs P | *Live individuals* | Pe | SR | 48.268 | 1 | **<0.001** |
| | | | T | 108.960 | 4 | **<0.001** |
| | | | SR × T | 35.189 | 4 | **<0.001** |
| | *Dead individuals* | Pe | SR | 0.063 | 1 | 0.801 |

**Table S11: Analysing the proportion of dead individuals within the first ~20 hours after coinfection and their Bt load upon death in selected regimes**. **(a)** The table provides a summary of Wilcoxon rank sum test on the proportion of dead individuals in each of the 4 replicate populations of C- vs M-beetles. **(b)** The table provides a summary of a generalised linear mixed model on log-transformed Bt load, fitted to a Gamma distribution, as a function of selection regime (SR) as fixed effect and replicate population (RP) as random effect [Model specification: Bacterial load ~ SR + (1|RP)].

| Experiment | Effect | P | | |
|---|---|---|---|---|
| **(a)** Proportion of dead individuals | SR | **0.0294** | | |
| | **Effect** | **$\chi^2$** | **df** | **P** |
| **(b)** Bt load | SR | 0.5094 | 1 | 0.4754 |

**Table S12:** The table provides a summary of a generalized linear model on log-transformed Bt load between 8 to 12 hours after coinfection (i.e., during the clearance phase of the Bt cells), fitted to a Gamma distribution [Model specification: Bacterial Load ~ Selection regime (SR) + Time points (T) + SR × T]. Significant effects are highlighted in bold.

| Effects | $\chi^2$ | df | P |
|---|---|---|---|
| SR | 0.5230 | 1 | 0.4695 |
| T | 4.3220 | 4 | **0.03** |
| SR × T | 0.9127 | 4 | 0.339 |

**Table S13:** List of differentially expressed genes with known immune-related functions, categorised as (a) pathogen receptors and immune receptors; (b) immune regulators; (c) inducible immune effectors, including antimicrobial peptides (AMPs) and lysozymes; fast-acting phenoloxidase mediated (d) melanisation response; and (e) production of reactive oxygen species.

| Gene name | Category | Functional role |
|---|---|---|
| PGRP SC1 a/b-like | Receptors | Short chain, soluble pathogen recognition receptor with scavenging function and phagocytosis (*15*) |
| PGRP SC2 | Receptors | Short chain, soluble pathogen recognition receptor that scavenges peptidoglycan components(*15*) |
| PGRP 2 | Receptors | Pathogen recognition receptor, activates Imd pathway (*16*) |
| PGRP LB | Receptors | Pathogen recognition receptor, prevents overactivation of Imd pathway (*15*) |
| PGRP LA | Receptors | Membrane bound pathogen recognition receptor (*15*) |
| PGRP LE | Receptors | Pathogen recognition receptor, functions in Imd pathway and autophagy (*15*) |
| GNBP 1 | Receptors | Pathogen recognition protein, activates Toll pathway (*15*) |
| GNPB 2 | Receptors | Pathogen recognition protein with uncharacterized function (*17*) |
| GNBP 3 | Receptors | Pathogen recognition protein, activates Toll pathway (*18*) |
| B-1,3-glucan binding protein 2 | Receptors | Directly binds to bacterial cells and helps in agglutination (*19*) |
| Integrin alpha subunit 2 | Receptors | Found in membranes of granulocytes , helps in phagocytosis and encapsulation (*20*) |
| TLR 3 | Receptors | Membrane bound receptor, helps in induction of AMPs (*21*) |
| Protein Toll | Receptors | Membrane bound receptor, helps in induction of AMPs (*21*) |
| Ankyrin 1 | Regulators | Negative regulator of Imd pathway (*22*) |
| Relish | Regulators | Positive regulator of Imd pathway (*23*) |
| Leukocyte elastase inhibitor | Regulators | Protect cells from proteases (*24*) |
| Decorin-like | Regulators | Damage associated molecular pattern, interacts with TLR (*25*) |
| Cactus | Regulators | Central regulator of Toll pathway signal transduction (*26*) |
| Serine protease inhibitor I/II | Regulators | Modulates multiple pathways involved in innate immunity (17) |
| Cathepsin L1 | Regulators | Cysteine protease regulates Toll and Imd pathway associated genes and autophagy (18) |
| Suppressor of cytokine signalling 2 | Regulators | Physiological regulators of cytokine responses (*27*) |

| Gene name | Category | Functional role |
|---|---|---|
| Pathogenesis related protein (664054) | Regulators | Belong to a functionally diverse group of proteins, that prevent pathogen colonisation (*28*) |
| Pathogenesis related protein (657457) | Regulators | Belong to a functionally diverse group of proteins, that prevent pathogen colonisation (*28*) |
| MIF homolog | Regulators | Proinflammatory cytokine, component of antimicrobial alarm system and stress response (*29*) |
| Serine protease persephone | Regulators | Detector of proteolytic activity and damage sensing, involved in Toll pathway (*30*) |
| LPS-induced TNF alpha homolog | Regulators | Cytokine involved in inflammation and immune signalling (23) |
| Insulin receptor substrate 1 | Regulators | Regulatory role in Insulin/Insulin-like signalling (*31*) |
| Apolipophorins I/II | Regulators | Immune stimulating protein found in hemolymph (*32*) |
| Attacin 1 | Effectors | Imd pathway specific AMP (2) |
| Attacin 2 | Effectors | Imd pathway specific AMP (2) |
| Coleoptericin | Effectors | Imd pathway specific AMP (2) |
| Coleoptericin-like | Effectors | Imd pathway specific AMP (2) |
| Tenecin 1 | Effectors | Imd pathway specific AMP (2) |
| Tenecin 2 | Effectors | Imd pathway specific AMP (2) |
| Tenecin 3 | Effectors | Imd/Toll pathway induced AMP(*33*) |
| Ctenidin 1 | Effectors | AMP active against gram-negative bacteria, have not been reported in *Tribolium castaneum* previously (*34*) |
| Holotricin 1/Defensin 3 | Effectors | Imd/Toll pathway induced AMP(*33*) |
| Cecropin 2 precursor | Effectors | Toll pathway specific AMP(*33*) |
| Lysozyme-like (100142316) | Effectors | Antibacterial enzyme that hydrolyzes peptidoglycan in bacterial cell wall (*35*) |
| Lysozyme-like (107398423) | Effectors | Antibacterial enzyme that hydrolyzes peptidoglycan in bacterial cell wall (*35*) |

| Gene name | Category | Functional role |
|---|---|---|
| Lysozyme precursor | Effectors | Antibacterial enzyme that hydrolyzes peptidoglycan in bacterial cell wall (*35*) |
| Hexamerin 1A/ Arylphorin precursor | PO response | Positively regulate hemolymph PPO (*36*) |
| Hexamerin 1B precursor | PO response | Positively regulate hemolymph PPO (*36*) |
| Hexamerin 2 | PO response | Positively regulate hemolymph PPO (*36*) |
| Serpin B6 (656375) | PO response | Regulation of prophenoloxidase activating proteinases (*37*) |
| Serpin B3 | PO response | Regulation of prophenoloxidase activating proteinases (*37*) |
| Serpin B12 | PO response | Regulation of prophenoloxidase activating proteinases (*37*) |
| Serpin B6 (658387) | PO response | Regulation of prophenoloxidase activating proteinases (*37*) |
| Laccase 2 | PO response | Oxidizes o-diphenols,p-diphenol and p-diamines (*38*) |
| Tyrosine decarboxylase | PO response | Conversion of Tyrosine to Tyramine (*39*) |
| Aspartate 1 decarboxylase | PO response | Converts Aspartate to Beta-alanine (*40*) |
| Phenoloxidase 2 | PO response | Key enzyme for conversion of Phenols to quinones that leads to melanin formation (*41*) |
| Pro-phenol oxidase subunit 2 | PO response | Catalyzes oxidation of phenols to quinones (41) |
| Yellow-g precursor | PO response | Helps in converting dopamine to dopamine-melanin (*42*) |
| Yellow g2-precursor | PO response | Helps in converting dopamine to dopamine-melanin (*42*) |
| GTP cyclohydrolase 1 | PO response | Essential for synthesis of tetrahydrobiopterin, a cofactor of TH (*43*) |
| Tyrosine hydroxylase (TH) | PO response | Converts tyrosine to L-DOPA (*42*) |
| DOPA decarboxylase | PO response | Coverts DOPA to dopamine (32) |
| Ebony | PO response | Catalyzes synthesis of precursor of yellow sclerotin N-Beta-alanyl dopamine (*42*) |

| Gene name | Category | Functional role |
| --- | --- | --- |
| Glutathione S-transferease 1 | ROS response | Quench reactive molecules to protect cell from oxidative damage (*44*) |
| Glutathione S-transferease | ROS response | Quench reactive molecules to protect cell from oxidative damage (*44*) |
| Peroxidase | ROS response | Protects cellular protein by detoxification of hydrogen peroxide (34) |
| Superoxide dismutase (656682) | ROS response | Catalyzes conversion of superoxides into oxygen and hydrogen peroxide, limiting the potential toxicity of ROS (34) |
| Superoxide dismutase (660957) | ROS response | Catalyzes conversion of superoxides into oxygen and hydrogen peroxide, limiting the potential toxicity of ROS (34) |
| Dual oxidase 1 | ROS response | Produces hydrogen peroxide by transferring electrons from intracellular NADPH to extracellular oxygen (*45*) |

**Table S14: Linear model fitting and gene expression profile of receptors**

Reduced model: Gene expression ~ Regime (R) + Infection Status (I) + Pathogen (P) + (Infection Status × Pathogen) + (Infection Status × Regime) + (Regime × Pathogen)

Type II MANOVA Tests: Wilk's test statistics

| Effects | Df | Test stat | F Statistics | Pr(>F) |
|---|---|---|---|---|
| R | 1 | 0.418 | 2.781 | **0.012** |
| I | 1 | 0.148 | 11.502 | **<0.001** |
| P | 2 | 0.014 | 15.089 | **<0.001** |
| I × P | 2 | 0.090 | 4.660 | **<0.001** |
| R × P | 2 | 0.298 | 1.661 | 0.059 |
| I × R | 1 | 0.555 | 1.605 | 0.147 |

Contrasts in gene expression profiles:

*B Line Tukey's HSD test statistics*

| Effects | Groups | Mean differences | Pr |
|---|---|---|---|
| Infection | C-Sham vs C-Bt | 0.424 | 0.472 |
| | B-Sham vs B-Bt | 0.455 | 0.414 |
| Selection | C-Sham vs B-Sham | -0.203 | 0.889 |
| | C-Bt vs B-Bt | -0.171 | 0.929 |

*M Line Tukey's HSD test statistics*

| Effects | Groups | Mean differences | Pr |
|---|---|---|---|
| Infection | C-Sham vs C-Mx | -0.605 | **0.002** |
| | M-Sham vs M-Mx | -0.555 | **0.004** |
| Selection | C-Sham vs M-Sham | -0.014 | 0.999 |
| | C-Mx vs M-Mx | -0.064 | 0.955 |

*P Line Tukey's HSD test statistics*

| Effects | Groups | Mean differences | Pr |
|---|---|---|---|
| Infection | C-Sham vs C-Pe | -0.781 | **0.001** |
| | P-Sham vs P-Pe | -0.786 | **0.001** |
| Selection | C-Sham vs P-Sham | -0.382 | 0.079 |
| | C-Pe vs P-Pe | -0.377 | 0.083 |

*For each pairwise comparison, the first alphabet denotes the regime followed by the infection treatment (sham infected or pathogen infected)

**Table S15: Linear model fitting and gene expression profile of regulators of immune pathways**

Reduced model: Gene expression ~ Infection Status (I) + Pathogen (P) + (Infection Status × Pathogen)

Type II MANOVA Tests: Wilk's test statistics

| Effects | Df | Test stat | F Statistics | Pr(>F) |
|---|---|---|---|---|
| I | 1 | 0.139 | 12.804 | **<0.001** |
| P | 2 | 0.016 | 14.292 | **<0.001** |
| I × P | 2 | 0.169 | 2.965 | **<0.001** |

Contrasts in gene expression profiles:

*B Line Tukey's HSD test statistics*

| Effects | Groups | Mean differences | Pr |
|---|---|---|---|
| Infection | C-Sham vs C-Bt | 0.178 | 0.814 |
| | B-Sham vs B-Bt | 0.341 | 0.371 |
| Selection | C-Sham vs B-Sham | -0.254 | 0.604 |
| | C-Bt vs B-Bt | -0.091 | 0.967 |

*M Line Tukey's HSD test statistics*

| Effects | Groups | Mean differences | Pr |
|---|---|---|---|
| Infection | C-Sham vs C-Mx | 0.232 | 0.298 |
| | M-Sham vs M-Mx | 0.120 | 0.774 |
| Selection | C-Sham vs M-Sham | 0.004 | 0.999 |
| | C-Mx vs M-Mx | 0.116 | 0.790 |

*P Line Tukey's HSD test statistics*

| Effects | Groups | Mean differences | Pr |
|---|---|---|---|
| Infection | C-Sham vs C-Pe | 0.583 | **<0.001** |
| | P-Sham vs P-Pe | 0.480 | **0.002** |
| Selection | C-Sham vs P-Sham | 0.265 | 0.083 |
| | C-Pe vs P-Pe | 0.369 | 0.013 |

*For each pairwise comparison, the first alphabet denotes the regime followed by the infection treatment (sham infected or pathogen infected)

**Table S16: Linear model fitting and gene expression profile of immune effectors including antimicrobial peptides**

Reduced model: Gene expression ~ Regime (R) + Infection Status (I) + Pathogen (P) + (Regime × Infection Status) + (Infection Status × Pathogen)

Type II MANOVA Tests: Wilk's test statistics

| Effects | Df | Test stat | F Statistics | Pr(>F) |
|---|---|---|---|---|
| R | 1 | 0.587 | 1.5118 | 0.174 |
| I | 1 | 0.157 | 11.604 | **<0.001** |
| P | 2 | 0.019 | 13.395 | **<0.001** |
| R × I | 1 | 0.461 | 2.521 | **0.020** |
| I × P | 2 | 0.069 | 6.044 | **<0.001** |

Contrasts in gene expression profiles:

*B Line Tukey's HSD test statistics*

| Effects | Groups | Mean differences | Pr |
|---|---|---|---|
| Infection | C-Sham vs C-Bt | -1.454 | **<0.001** |
| | B-Sham vs B-Bt | -1.224 | **0.001** |
| Selection | C-Sham vs B-Sham | 0.110 | 0.9662 |
| | C-Bt vs B-Bt | 0.3405 | 0.514 |

*M Line Tukey's HSD test statistics*

| Effects | Groups | Mean differences | Pr |
|---|---|---|---|
| Infection | C-Sham vs C-Mx | 0.6711 | **0.001** |
| | M-Sham vs M-Mx | 0.256 | 0.266 |
| Selection | C-Sham vs M-Sham | -0.062 | 0.964 |
| | C-Mx vs M-Mx | 0.353 | 0.084 |

*P Line Tukey's HSD test statistics*

| Effects | Groups | Mean differences | Pr |
|---|---|---|---|
| Infection | C-Sham vs C-Pe | 0.3179 | 0.093 |
| | P-Sham vs P-Pe | 0.6483 | **<0.001** |
| Selection | C-Sham vs P-Sham | 0.1695 | 0.5304 |
| | C-Pe vs P-Pe | -0.1608 | 0.5715 |

*For each pairwise comparison, the first alphabet denotes the regime followed by the infection treatment (sham infected or pathogen infected)

**Table S17: Linear model fitting and gene expression profile of phenoloxidase response related components**

Reduced model: Gene expression ~ Infection Status (I) + Pathogen (P)+ (Infection Status × Pathogen)

Type II MANOVA Tests: Wilk's test statistics

| Effects | Df | Test stat | F Statistics | Pr(>F) |
|---|---|---|---|---|
| I | 1 | 0.106 | 10.595 | **<0.001** |
| P | 2 | 0.026 | 6.497 | **<0.001** |
| I × P | 2 | 0.137 | 2.138 | **<0.001** |

Contrasts in gene expression profiles:

*B Line Tukey 's HSD test statistics*

| Effects | Groups | Mean differences | Pr |
|---|---|---|---|
| Infection | C-Sham vs C-Bt | -1.509 | **<0.001** |
| | B-Sham vs B-Bt | -0.790 | **<0.001** |
| Selection | C-Sham vs B-Sham | -0.227 | 0.269 |
| | C-Bt vs B-Bt | 0.491 | 0.006 |

*M Line Tukey's HSD test statistics*

| Effects | Groups | Mean differences | Pr |
|---|---|---|---|
| Infection | C-Sham vs C-Mx | -0.538 | **0.03** |
| | M-Sham vs M-Mx | -0.652 | **0.009** |
| Selection | C-Sham vs M-Sham | 0.189 | 0.672 |
| | C-Mx vs M-Mx | 0.303 | 0.306 |

*P Line Tukey's HSD test statistics*

| Effects | Groups | Mean differences | Pr |
|---|---|---|---|
| Infection | C-Sham vs C-Pe | 0.023 | 0.9973 |
| | P-Sham vs P-Pe | -0.877 | **<0.001** |
| Selection | C-Sham vs P-Sham | -0.321 | 0.099 |
| | C-Pe vs P-Pe | 0.581 | **0.002** |

*****For each pairwise comparison, the first alphabet denotes the regime followed by the infection treatment (sham infected or pathogen infected)

**Table S18: Linear model fitting and gene expression profile of reactive oxygen species response molecules**

Reduced model: Gene expression ~ Infection Status (I) + Pathogen (P) + (Infection Status × Pathogen)

Type II MANOVA Tests: Wilk's test statistics

| Effects | Df | Test stat | F Statistics | Pr(>F) |
|---|---|---|---|---|
| I | 1 | 0.226 | 21.082 | **<0.001** |
| P | 2 | 0.179 | 8.403 | **<0.001** |
| I × P | 2 | 0.543 | 2.198 | **0.020** |

Contrasts in gene expression profiles:

*B Line Tukey's HSD test statistics*

| Effects | Groups | Mean differences | Pr |
|---|---|---|---|
| Infection | C-Sham vs C-Bt | -0.529 | 0.079 |
| | B-Sham vs B-Bt | -0.782 | **0.008** |
| Selection | C-Sham vs B-Sham | 0.323 | 0.390 |
| | C-Bt vs B-Bt | 0.071 | 0.982 |

*M Line Tukey's HSD test statistics*

| Effects | Groups | Mean differences | Pr |
|---|---|---|---|
| Infection | C-Sham vs C-Mx | -1.242 | **0.009** |
| | M-Sham vs M-Mx | -1.413 | **0.003** |
| Selection | C-Sham vs M-Sham | 0.111 | 0.984 |
| | C-Mx vs M-Mx | 0.282 | 0.806 |

*P Line Tukey's HSD test statistics*

| Effects | Groups | Mean differences | Pr |
|---|---|---|---|
| Infection | C-Sham vs C-Pe | -1.973 | **<0.001** |
| | P-Sham vs P-Pe | -1.847 | **<0.001** |
| Selection | C-Sham vs P-Sham | -0.266 | 0.845 |
| | C-Pe vs P-Pe | -0.392 | 0.636 |

*****For each pairwise comparison, the first alphabet denotes the regime followed by the infection treatment (sham infected or pathogen infected)

**Table S19: Canonical correlation between the expression profile of receptors and phenotypic profile (Bacterial load and hazard ratio)**

Canonical correlation statistics:

| | Canonical coefficients | |
|---|---|---|
| | **Dimension 1 (Corr: 0.84)** | **Dimension 2 (Corr: 0.64)** |
| **Gene expression profile (LDA)** | | |
| LD1 | -0.116 | 0.039 |
| LD2 | 0.262 | -0.471 |
| LD3 | -0.197 | -0.259 |
| **Phenotypic profile** | | |
| Bacterial load | -0.929 | -0.302 |
| Hazard ratio | 0.025 | 0.101 |

Linear regression statistics:

| Selection regime | Residual S.E. | Adj. $R^2$ | F statistics | df | P value |
|---|---|---|---|---|---|
| B regime | 0.49 | 0.18 | 2.55 | 6 | 0.161 |
| M regime | 0.27 | 0.47 | 7.18 | 6 | **0.036** |
| P regime | 0.38 | 0.77 | 24.14 | 6 | **0.003** |

**Table S20: Canonical correlation between the expression profile of regulators of immune pathways and phenotypic profile (Bacterial load and hazard ratio)**

Canonical correlation statistics:

| | Canonical coefficients | |
|---|---|---|
| | **Dimension 1 (Corr: 0.72)** | **Dimension 2 (Corr: 0.67)** |
| **Gene expression profile (LDA)** | | |
| LD1 | -0.166 | -0.045 |
| LD2 | 0.444 | -0.673 |
| LD3 | 0.499 | 0.123 |
| **Phenotypic profile** | | |
| Bacterial load | 0.687 | 0.695 |
| Hazard ratio | 0.024 | -0.101 |

Linear regression statistics:

| Selection regime | Residual S.E. | Adj. $R^2$ | F statistics | df | P value |
|---|---|---|---|---|---|
| B regime | 0.44 | 0.77 | 24.69 | 6 | **0.002** |
| M regime | 0.75 | -0.14 | 0.13 | 6 | 0.732 |
| P regime | 0.72 | 0.54 | 9.13 | 6 | **0.023** |

**Table S21: Canonical correlation between the expression profile of immune effectors and phenotypic profile (Bacterial load and hazard ratio)**

Canonical correlation statistics:

| | Canonical coefficients | |
|---|---|---|
| | Dimension 1 (Corr: 0.68) | Dimension 2 (Corr: 0.60) |
| **Gene expression profile (LDA)** | | |
| LD1 | -0.058 | -0.063 |
| LD2 | 0.115 | -0.599 |
| LD3 | -0.410 | 0.270 |
| **Phenotypic profile** | | |
| Bacterial load | 0.859 | 0.466 |
| Hazard ratio | -0.006 | -0.104 |

Linear regression statistics:

| Selection regime | Residual S.E. | Adj. $R^2$ | F statistics | df | P value |
|---|---|---|---|---|---|
| B regime | 0.33 | 0.17 | 2.46 | 6 | 0.167 |
| M regime | 0.48 | 0.41 | 5.91 | 6 | 0.05 |
| P regime | 0.61 | 0.50 | 7.90 | 6 | **0.031** |

**Table S22: Canonical correlation between the expression profile of phenoloxidase response molecules and phenotypic profile (Bacterial load and hazard ratio)**

Canonical correlation statistics:

| | Canonical coefficients | |
|---|---|---|
| | Dimension 1 (Corr: 0.77) | Dimension 2 (Corr: 0.21) |
| **Gene expression profile (LDA)** | | |
| LD1 | 0.218 | -0.110 |
| LD2 | 0.109 | 0.382 |
| LD3 | 0.516 | 0.355 |
| **Phenotypic profile** | | |
| Bacterial load | -0.913 | 0.349 |
| Hazard ratio | 0.020 | -0.102 |

Linear regression statistics:

| Selection regime | Residual S.E. | Adj. $R^2$ | F statistics | df | P value |
|---|---|---|---|---|---|
| B regime | 0.60 | 0.46 | 6.98 | 6 | **0.038** |
| M regime | 0.35 | 0.29 | 3.80 | 6 | 0.099 |
| P regime | 0.63 | 0.14 | 2.19 | 6 | 0.189 |

**Table S23: Canonical correlation between the expression profile of ROS response and phenotypic profile (Bacterial load and hazard ratio)**

Canonical correlation statistics:

| | Canonical coefficients | |
|---|---|---|
| | Dimension 1 (Corr: 0.63) | Dimension 2 (Corr: 0.17) |
| **Gene expression profile (LDA)** | | |
| LD1 | -0.212 | -0.405 |
| LD2 | 0.418 | -0.233 |
| LD3 | 0.388 | -0.714 |
| **Phenotypic profile** | | |
| Bacterial load | -0.834 | -0.509 |
| Median lifespan | 0.001 | 0.104 |

Linear regression statistics:

| Selection regime | Residual S.E. | Adj. $R^2$ | F statistics | df | P value |
|---|---|---|---|---|---|
| B regime | 0.53 | 0.22 | 2.95 | 6 | 0.136 |
| M regime | 0.45 | -0.05 | 0.66 | 6 | 0.446 |
| P regime | 1.03 | 0.28 | 3.72 | 6 | 0.102 |

**Table S24: Least square regression statistics between expression values of immune molecule categories and hazard ratio.**

| Immune categories | Selection regime | Residual S.E. | Adj. $R^2$ | F-statistics | df | P value |
|---|---|---|---|---|---|---|
| Receptor | B regime | 0.35 | -0.13 | 0.17 | 6 | 0.690 |
| | M regime | 0.12 | 0.05 | 1.38 | 6 | 0.284 |
| | P regime | 0.16 | 0.60 | 11.58 | 6 | **0.014** |
| Regulator | B regime | 0.12 | 0.17 | 2.46 | 6 | 0.168 |
| | M regime | 0.12 | 0.47 | 7.14 | 6 | **0.037** |
| | P regime | 0.10 | -0.003 | 0.98 | 6 | 0.361 |
| Effector | B regime | 0.24 | 0.31 | 4.12 | 6 | 0.089 |
| | M regime | 0.20 | 0.15 | 2.28 | 6 | 0.182 |
| | P regime | 0.15 | -0.12 | 0.25 | 6 | 0.635 |
| PO response | B regime | 0.20 | 0.59 | 11.22 | 6 | 0.015 |
| | M regime | 0.21 | 0.18 | 2.55 | 6 | 0.161 |
| | P regime | 0.18 | 0.66 | 14.73 | 6 | **0.009** |
| ROS response | B regime | 0.17 | -0.12 | 0.28 | 6 | 0.617 |
| | M regime | 0.57 | -0.17 | 0.001 | 6 | 0.945 |
| | P regime | 0.49 | 0.19 | 2.62 | 6 | 0.157 |

**Table S25: Least square regression statistics between expression values of immune molecule categories and bacterial load.**

| Immune categories | Selection regime | Residual S.E. | Adj. $R^2$ | F-statistics | df | P value |
|---|---|---|---|---|---|---|
| Receptor | B regime | 0.35 | -0.13 | 0.21 | 6 | 0.662 |
| | M regime | 0.13 | -0.16 | 0.05 | 6 | 0.828 |
| | P regime | 0.14 | 0.70 | 17.36 | 6 | **0.006** |
| Regulator | B regime | 0.15 | -0.17 | 0.0003 | 6 | 0.986 |
| | M regime | 0.09 | 0.70 | 17.52 | 6 | **0.006** |
| | P regime | 0.10 | -0.02 | 0.89 | 6 | 0.382 |
| Effector | B regime | 0.31 | -0.16 | 0.02 | 6 | 0.902 |
| | M regime | 0.21 | -0.01 | 0.93 | 6 | 0.371 |
| | P regime | 0.15 | -0.16 | 0.04 | 6 | 0.851 |
| PO response | B regime | 0.30 | 0.05 | 1.40 | 6 | 0.282 |
| | M regime | 0.16 | 0.54 | 9.11 | 6 | **0.023** |
| | P regime | 0.19 | 0.65 | 13.96 | 6 | **0.010** |
| ROS response | B regime | 0.14 | 0.22 | 3.02 | 6 | 0.133 |
| | M regime | 0.52 | 0.02 | 1.17 | 6 | 0.322 |
| | P regime | 0.48 | 0.23 | 3.05 | 6 | 0.131 |

**Table S26: Analysing fecundity across infection treatments in the baseline stock population**. The table provides a summary of a generalised linear model on the data describing the total number of eggs laid by each female, fitted to a Poisson distribution, as a function of infection treatments (I) (e.g., sham-infection, infection with Bt, Pe and Mx) [Model specification: Egg count ~ Infection treatment (I)], **(a)** between 6–42 hours post-infection (hpi) **(b)** between 78–114hpi. Pairwise contrasts between different treatments are based on Tukey's HSD. Significant differences are highlighted in bold.

| **Infection phase** | **Effects** | $\chi^2$ | **df** | **P** |
|---|---|---|---|---|
| **(a)** 6-42hpi | I | 44.02 | 3 | **<0.001** |
| **(b)** 78-114hpi | I | 149.76 | 3 | **<0.001** |

Pairwise comparison

| **Infection phase** | **Contrasts** | **P** |
|---|---|---|
| (a) 6-42hpi | Sham vs Bt | **<0.001** |
| | Sham vs Pe | **<0.001** |
| | Sham vs Mx | **<0.001** |
| | Bt vs Pe | 0.134 |
| | Bt vs Mx | 0.997 |
| | Mx vs Pe | 0.121 |
| (b) 78-114hpi | Sham vs Bt | 0.988 |
| | Sham vs Pe | **<0.001** |
| | Sham vs Mx | **<0.001** |
| | Bt vs Pe | **<0.001** |
| | Bt vs Mx | **<0.001** |
| | Mx vs Pe | **<0.001** |

**Table S27: Phenoloxidase activity after Bt infection in B and C beetles.** The table provides a summary of a Wilcoxon rank sum test on phenoloxidase activity after Bt-infection (I) or sham infection (S) in replicate populations 1 and 2 of B and C-regimes. Significant differences are highlighted in bold.

| **Replicate population** | **Regime** | **Contrasts** | **P** |
|---|---|---|---|
| 1 | C | I vs S | **0.01** |
| | B | I vs S | 0.123 |
| 2 | C | I vs S | **0.01** |
| | B | I vs S | 0.143 |
